# A Reinforcement Learning and Sequential Sampling Model Constrained by Gaze Data

**DOI:** 10.1101/2025.08.27.672620

**Authors:** William M. Hayes, Melanie J. Touchard

**Affiliations:** Psychology Department, Binghamton University State University of New York, Binghamton, NY, 13902

## Abstract

Reinforcement learning models can be combined with sequential sampling models to fit choice-RT data. The combined models, known as RL-SSMs, explain a wide range of choice-RT patterns in repeated decision tasks. The present study shows how constraining an RL-SSM with eye gaze data can further enhance its predictive ability. Our model assumes that learned option values and relative gaze independently influence the accumulation of evidence prior to choice. We evaluated the model on data from two eye-tracking experiments (total N = 133) and find that it makes better out-of-sample predictions than other models with different ways of integrating values and gaze at the decision stage. Further, we show that it captures a variety of empirical effects, including the finding that choices become more accurate as the higher-value option receives a greater proportion of the total fixation time. The model can be used to understand how learned option values interact with visual attention to influence choice, joining together two major—but mostly separate—research traditions in the cognitive science of decision making.

## INTRODUCTION

Whether deciding what to eat for lunch, which route to take to avoid traffic, or who to trust for financial advice, many of our decisions require us to rely on information learned through experience. Our preferences in these situations are often influenced by the outcomes of previous choices. However, several other factors can influence these decisions, including the time spent deliberating and the amount of attention allocated to the different alternatives at the time of choice ^1,2^.

Traditionally, decisions from experience have been studied using reinforcement learning models (RL), which describe how decision makers adjust their choice behavior in response to feedback from the environment to maximize reward ^3^. Most RL models use a static choice rule (softmax) to select actions based on expected values ^4^. This choice rule has key limitations. First, it does not account for choice response times (RTs), which are known to reflect decision difficulty and speed-accuracy tradeoffs ^5,6^. For example, it may take longer to make a decision if the available options are subjectively similar, or if the decision maker wants to find the best possible option as opposed to one that is just good enough. Second, in its usual form, the static choice rule in RL models does not account for attentional biases in action selection ^7^ (but see ^8,9^). A decision maker may choose a less desirable option (based on their previous experience) simply because they spend more time fixating on it. Such effects would be missed by standard RL models.

Meanwhile, separate lines of research on perceptual and value-based decision making have led to the development of sequential sampling models (SSMs) that can simultaneously account for choices and RTs ^10–13^, as well as attentional biases ^14,15^. Though exact implementations vary, SSMs generally assume that choices occur through a dynamic process in which evidence (or preference) for choosing a particular option accumulates over time until a decision threshold is reached. The time taken to reach the threshold, plus a small amount of time for stimulus encoding and response execution, determines the RT. In value-based decisions, the speed of evidence accumulation, or drift rate, for a particular option reflects its subjective value to the decision maker (or its relative subjective value). The height of the decision threshold reflects the decision maker’s response caution, or the amount of evidence they require to make a choice.

SSMs have also been used to model the role of overt visual attention, or gaze, in value-based decisions ^16,17^. This line of work has demonstrated robust gaze biases. In particular, the longer an option is fixated, the more likely it is to be chosen ^14,15,18–21^. This effect persists after controlling for the subjective values of the choice options, and therefore cannot be explained away as merely a tendency to look longer at high-value options. Rather, gaze appears to either amplify ^22^ or add to ^19,23^ the value of an option during evidence accumulation. For example, in the attentional drift diffusion model (aDDM; ^14^), looking at an option temporarily biases the drift rate toward the fixated option by discounting the value of the unfixated option: a value-amplifying effect. Other models assume that gaze adds a fixed bonus to the drift rate that is independent of the option value ^19^. Which mechanism is best—additive or multiplicative— may depend on the choice task ^22^.

In summary, while RL models can explain how preferences are learned through trial-and-error, SSMs give a more complete account of the decision stage, incorporating choice RTs and (optionally) gaze biases. Fortunately, the two approaches can be combined by substituting an SSM for the static choice rule in an RL model ^24–29^; for a review, see ^1^. In this approach, learning occurs over trials according to an incremental RL mechanism, and choices are made through a sequential sampling process, with the drift rate(s) depending on the current learned values for each option. These combined RL-SSMs explain a variety of behavioral patterns in decisions from experience, including the effects of learning and choice difficulty on accuracy and response time ^26^. However, to our knowledge, there is currently no RL-SSM that accounts for gaze biases and how they interact with value learning to influence behavior.

In the present study, we introduce an RL-SSM that can account for trial-and-error learning, choice RTs, and attentional biases in decisions from experience. The model explains a variety of empirical effects: Increasing accuracy and faster RTs over the course of learning, individual differences in overall accuracy and decision speed, and (in Experiment 2) context-dependent valuation ^30–33^. Unlike existing RL-SSMs, it also captures gaze effects: The longer an option is fixated relative to other options, the more likely it is to be chosen. The model has several practical advantages: it is lightweight, analytically tractable, and readily scalable to more complex tasks. In the following, we briefly describe the experiments that were used to evaluate the model before turning to a more detailed description of the model itself.

## RESULTS

### Overview of Experiments

We evaluated our model on data from two eye-tracking experiments (Figure 1A). Although both experiments were originally designed to examine context-dependent valuation in RL, the resulting data sets provide a rich testbed for evaluating RL-SSMs.

**Figure 1:**
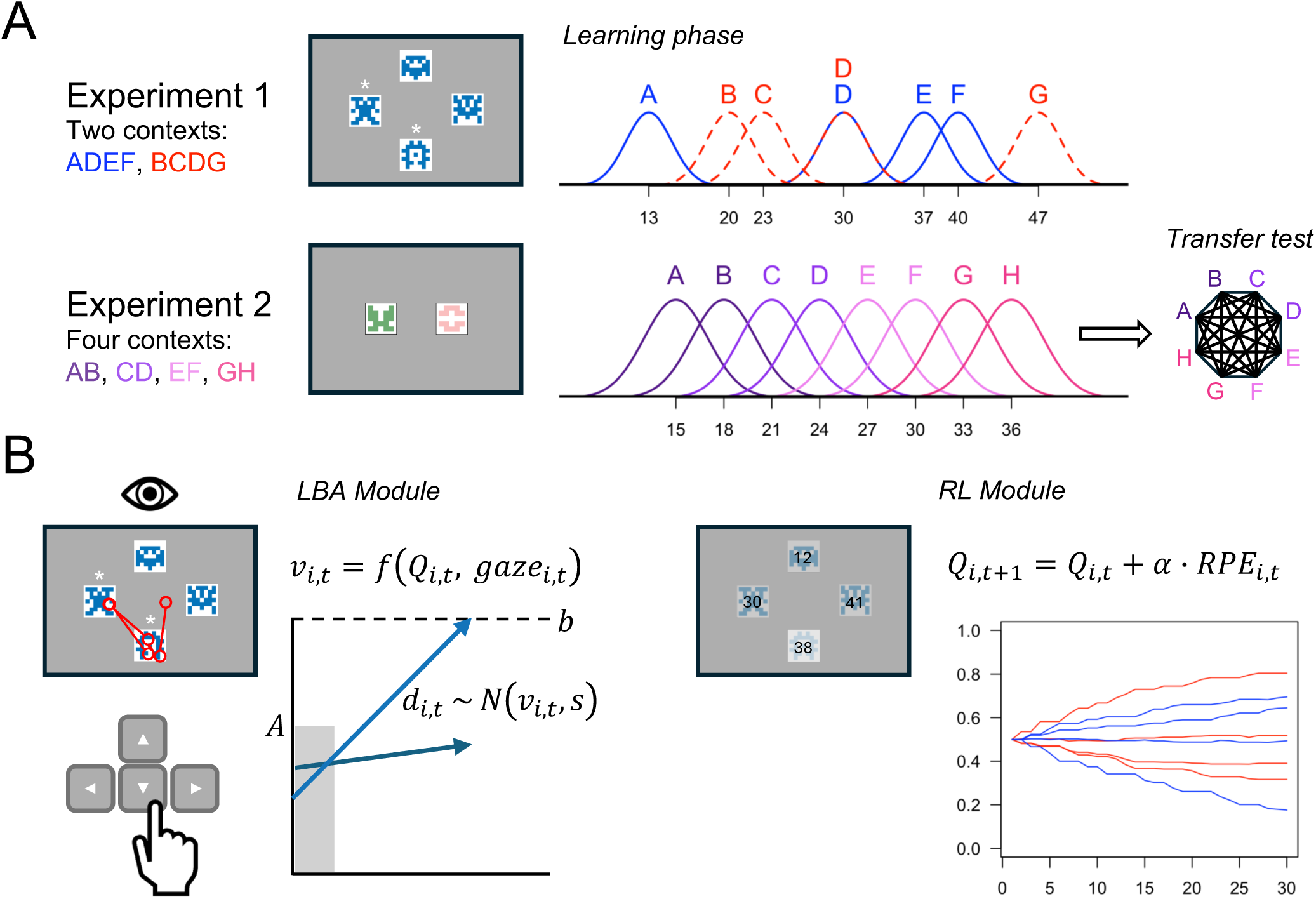
Overview of experiments and model. (A) In the learning phase, symbols were encountered in fixed groupings (contexts). On every trial, the symbols from a randomly selected context were presented on screen. In Experiment 1, only two of the four symbols were available to choose (indicated with asterisks). The available symbols varied from trial to trial. Choices were followed by probabilistic reward feedback from all options seen on that trial. Experiment 2 included a transfer test after the learning phase that involved choices between all possible pairs of symbols without feedback. The outcome distributions in both experiments were Gaussian with option-specific means (SD = 2). (B) The LBA module uses linear ballistic accumulation to make decisions. The mean drift rates (*v_i,t_*) are assumed to be a function of the current estimated values of the available symbols (*Q_i,t_*) and the proportional gaze allocated to each symbol during the time interval between trial onset and choice (*gaze_i,t_*). The RL module updates the estimated value of each symbol in response to feedback using a prediction error-driven learning rule (RPE = reward prediction error).

The task in the first experiment (N = 83) was adapted from a prior study ^31^. Participants encountered several choices between eight symbols grouped in two sets of four. The symbol groupings, or contexts, remained fixed across the 60 learning phase trials. Every trial began with the presentation of the four symbols from a randomly selected context, but only two of the four were available to choose (indicated by asterisks). After making a choice, participants received probabilistic reward feedback from all four symbols, including the ones that they could not select. The rewards were point values drawn from Gaussian distributions with different expected values (EVs) and rounded to whole numbers. The instructed goal was to learn which symbols had the highest value and to maximize the total number of points earned.

The task in the second experiment (N = 50) was adapted from another prior study ^34^, and parts of the data have been reported elsewhere ^35^. In this experiment, the symbols were grouped in four 2-option contexts instead of two 4-option contexts, the learning phase was longer, and the task ended with a transfer test in which participants encountered choices between all possible pairs of symbols without receiving feedback. The transfer test was designed to reveal whether participants had learned the absolute EVs of the symbols (context-independent) or the relative values (context-dependent). In the latter case, we should observe a predictable pattern of suboptimal preferences in the transfer test. For example, given a choice between options B and C, absolute values would dictate choosing C because it has a higher absolute EV (21 compared to 18). Relative values, in contrast, would dictate choosing B because it was better locally in its original context.

The trials were self-paced in both experiments, which allowed us to measure choice RTs. Importantly, because the choice contexts were presented in a random order, participants could not anticipate which set of symbols they would see on the next trial and decide beforehand which symbol to choose ^9,36^. This is crucial for our modeling assumption that gaze directly biases the evidence accumulation process leading up to choice. Next, we introduce the gaze-constrained RL-SSM that we used to fit choice-RT data in both experiments.

### Computational Model

Our computational model combines a linear ballistic accumulation (LBA) module for decision making ^13^ and an RL module for learning the values of options through experience (Figure 1B). The RL module maintains estimated values for each option that it updates in response to feedback from the environment. At choice time, the current value estimates for the available options are mapped to the drift rates in the LBA module, which determine how the evidence accumulation process unfolds. The model assumes that drift rates are also influenced by trial-to-trial fluctuations in visual attention to the available options, which allows it to explain a wider range of behavioral effects compared to previous RL-SSMs ^25–28^.

The LBA module assumes that choices result from a dynamic sequential sampling process. Evidence (or preference) accumulates in a linear trajectory toward the decision threshold (*b*), separately for each of the available options. The chosen option is the one whose accumulator reaches the threshold first. The time taken for the winning accumulator to reach the threshold, plus a small amount of non-decision time (*t*_0_), determines the choice RT. There are two sources of between-trial variability in the LBA module. First, the starting point for each option’s accumulator is assumed to be drawn independently on each trial from a Uniform(0*, A*) distribution. Second, the drift rate, or slope, of the *i*th option’s accumulator on trial *t*, *d_i,t_*, is assumed to be drawn independently from a Normal(*v_i,t_, s*) distribution. In our model, the mean of the drift rate distribution for the *i*th option on trial *t* depends on two inputs: the RL module’s current estimate of the option’s value, *Q_i,t_*, and 2) the relative gaze received by the option in the current trial. That is, we define a *linking function* ^1^ that maps learned Q-values and gaze onto the mean drift rates:

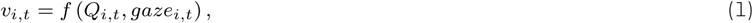

where *gaze_i,t_* is defined as the proportional amount of time that option *i* is fixated in trial *t* during the time period from trial onset to choice (see *Gaze Effects*). The linking function is what connects the RL and LBA modules together.

In our model, the mean drift rate for the *i*th option on trial *t* is computed as:

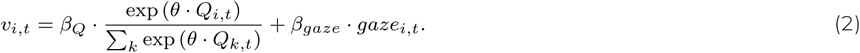

The right-hand side of this equation consists of two terms: The first is the softmax function applied to *Q_i,t_* with inverse temperature *θ*, and the second is the proportional gaze for option *i* on trial *t*. The scaling parameters *β_Q_* and *β_gaze_* represent the relative contributions of learned values and current gaze allocation, respectively, on the mean drift rates. They also serve to scale the drift rates appropriately. Following ^28^, we fixed *θ* = 50 in all model fits to reduce the number of free parameters and improve identifiability.

Two points are worth noting about the form of the linking function (Eq. 2). First, our use of the softmax is based on prior work demonstrating that a non-linear mapping from Q-values to drift rates improves the fit of RL-SSMs ^20,26,28,29^. Models that use a linear mapping tend to underestimate choice accuracy for difficult choices between options with similar Q-values ^26^. Because the softmax function with *θ* = 50 is very sensitive to small differences in the underlying Q-values, it produces sharper decision boundaries and thus greater overall accuracy (Fig. S1). The softmax also causes RTs to depend only on the difference between Q-values, with longer RTs for difficult choices (i.e., small Q-value differences). Linear versions of the model instead predict a magnitude effect: RTs are longer when both Q-values are small and faster when both Q-values are large (Fig. S1).

Second, our model assumes that relative gaze has an additive effect on drift rates that is independent of learned option values: As an option receives a greater proportion of the total fixation time leading up to a choice, the likelihood that it will be chosen increases, regardless of its value (Fig. S2). In contrast, models such as the aDDM assume a multiplicative effect in which gaze amplifies the value of the fixated option ^14^. According to this account, gaze will have a stronger effect on choice for higher-value options (see the “Q * gaze” model in Fig. S2). Although there is considerable evidence for a multiplicative gaze effect, the evidence from RL tasks is less clear ^22^. One study that examined gaze effects in the test phase of an RL task (i.e., after learning had taken place) found support for an additive gaze effect ^19^. In another RL study, a large subset of participants exhibited choice behavior that was inconsistent with core predictions of the aDDM ^36^; however, an additive model was not explicitly considered. It is difficult to directly compare our model to the aDDM due to differences in the underlying mathematics (e.g., racing accumulators versus drift diffusion ^37^); however, we did test alternate versions of our model that assume a multiplicative gaze effect. The overall evidence from both experiments favored the additive gaze model in Eq. 2 (see *Model Comparison* section). We therefore retained the additive assumption in our final model and present fits from the multiplicative models in the Supplemental Material.

Once a choice is made and feedback is presented, the RL module updates the expected values for each option using a standard delta learning rule ^38^:

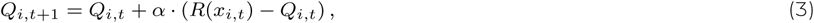

where *Q_i,t_*_+1_ is the updated expected value for the *i*th option, *Q_i,t_* is the previous expected value, and *α* is the learning rate parameter. The goal of this learning rule is to minimize the reward prediction error, *R*(*x_i,t_*)−*Q_i,t_*, or the difference between observed and expected outcomes for each option. Based on a growing body of research on context-dependent valuation in RL ^30–32^, our model assumes that the subjective value of *x_i,t_*, the outcome from option *i* on trial *t*, is a weighted combination of two range-normalized values:

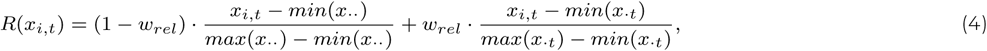

where the first term uses the minimum and maximum outcomes across all trials, and the second term uses the minimum and maximum outcomes on the current trial alone. Consider Option B in the task from Experiment 2 (Figure 1A). Option B’s outcomes are low with respect to the global range of outcomes across all contexts; thus, it will produce lower values for the first range-normalized term. However, it produces larger outcomes than Option A most of the time, and because these two options are always presented together, Option B will frequently have high values for the second range-normalized term (i.e., 1.0). As the relative encoding parameter *w_rel_* approaches 1, the second term will dominate and options will be subjectively valued based on their *relative* rank within the local context. As *w_rel_* approaches 0, the first term will dominate and subjective values will reflect global ranks (context-independent). The transfer test in Experiment 2 is critical for estimating *w_rel_*, as it pits options whose values were learned in one context against options whose values were learned in a different context. Because there was no transfer test in Experiment 1, for that experiment we fixed *w_rel_* = 0 (absolute encoding).

To summarize, our model includes seven free parameters: learning rate *α*, relative encoding *w_rel_*, drift scaling parameters *β_Q_*and *β_gaze_*, start point upper bound *A*, decision threshold *b*, and non-decision time *t*_0_. The first two control the RL module and the last five control the LBA module. Several plausible models can be constructed that differ in the way that Q-values and gaze are mapped onto the mean drift rates (Eq. 1). As discussed in the next section, the model that we ultimately selected was superior to others at making one-step-ahead predictions. Later, we will show that it also captures the key behavioral effects in our data.

### Model Comparison

A model should not only fit the training data well, but should also generalize to unseen data from the same data-generating process ^39^. With this in mind, we compared a set of candidate models using a metric known as accumulative one-step-ahead prediction error (APE) ^40^. APE is a measure of out-of-sample prediction accuracy appropriate for time series data. It measures a model’s ability to predict the next observation in a sequence, *y_n_*_+1_, given only the previous observations, *y*_1_*, …, y_n_*. Unlike similar metrics such as leave-one-out cross validation, APE only uses past observations to predict the future; it never uses future observations to predict the past. This makes it well-suited for RL problems. For a discussion of its other advantages and connections to other approaches, see ^40^. Formally, the APE with logarithmic loss for model *M_j_* is defined as

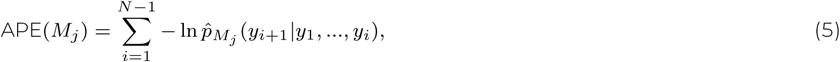

where ln *p*^*_Mj_* (*y_i_*_+1_|*y*_1_*, …, y_i_*) represents the log-likelihood that model *M_j_* assigns to observation *i* + 1, given the first *i* observations. APE is straightforward to implement, albeit computationally expensive: For an individual data set with *N* observations (i.e., choice-RT pairs), we fit the model *N* − 1 times, each time fitting a larger subset of the data. On each iteration, we use the fitted parameters to compute the log-likelihood of the next unseen observation. Then, the one-step-ahead log-likelihoods are summed across observations. The model with the lowest APE is taken as the one that generalizes best to unseen data. In practice, it may not be possible to estimate the model parameters precisely until the model is fit to a minimum amount of data ^40^. For the present application, we set the minimum number of data points to *n* = 5 (i.e., the sum in Eq. 5 is from *i* = 5 to *N* − 1).

We compared the model described in the previous section against six others, which differed in 1) the linking function (linear or non-linear/softmax), and 2) the type of gaze effect (none, multiplicative, or additive). For the models with a non-linear linking function, we tested two forms of multiplicative gaze effects: One in which gaze modulates the Q-values prior to the softmax transformation, and another in which gaze modulates the output of the transformation. Full descriptions of all models are provided in Appendix A. As shown in Figure 2A, the model with 1) a non-linear (softmax) linking function and 2) an additive gaze effect had the lowest mean APE in both experiments. Moreover, its advantage over the other models was significant (one-sample t-tests on APE difference scores [*H*_0_ : *µ* = 0]; Exp. 1: ps < .004; Exp. 2: ps < .022). We conclude that this particular model generalizes better to unseen data in the context of our experimental tasks.

**Figure 2:**
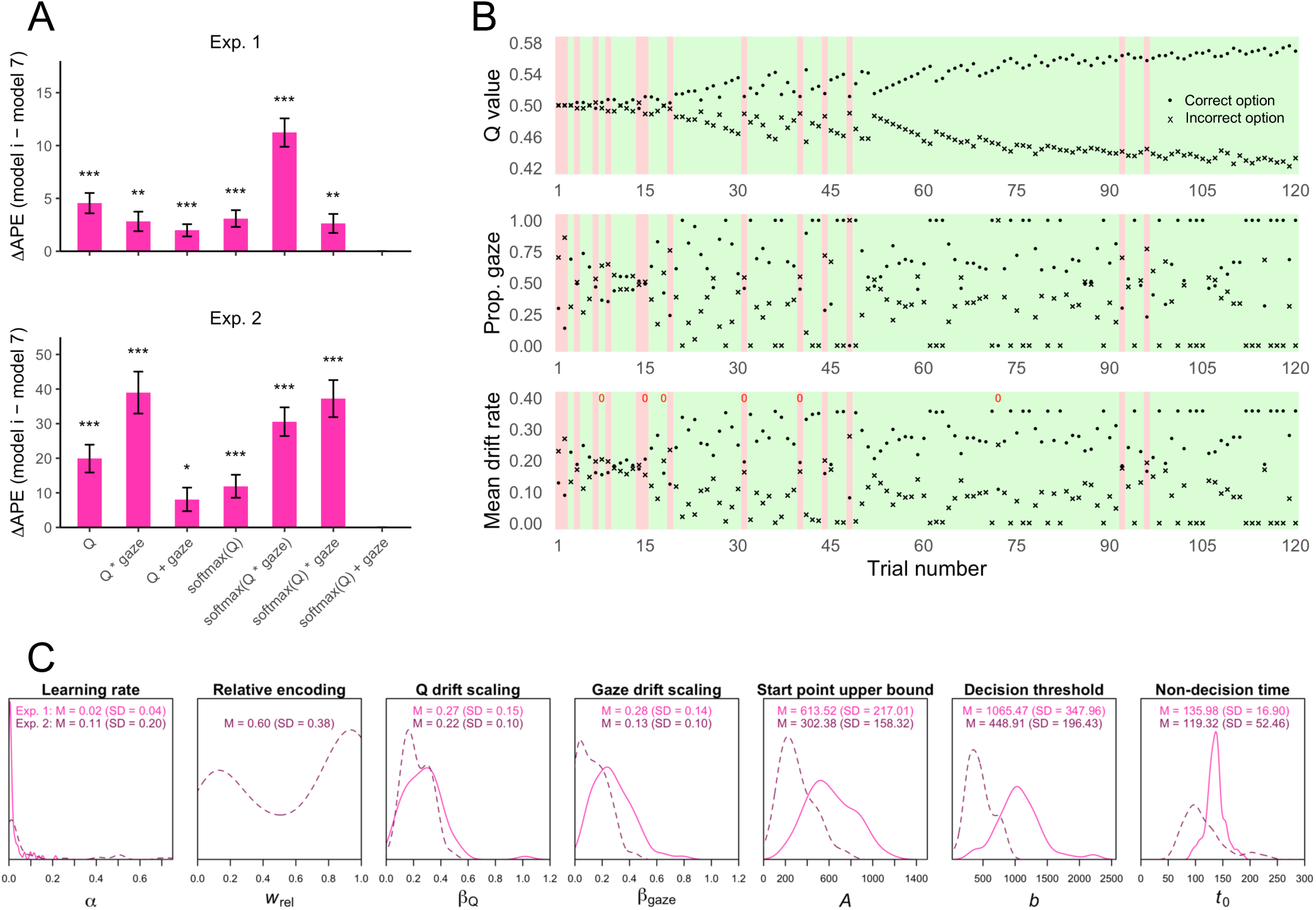
Model comparison results, latent variables, and estimated parameters. (A) Each bar shows the mean difference in accumulative one-step-ahead prediction error (APE; Eq. 5) between one of the alternative models and the winning model, softmax(Q) + gaze, averaging across participants. In all comparisons, the mean difference in APE was significantly above zero, indicating that the winning model generalized better to unseen data. *p < .05, **p < .01, ***p < .001 (one-sample t-tests). (B) Trial-to-trial Q-values, proportional gaze data, and mean drift rates for participant 32 in Experiment 2 (parameter estimates: *α* = .008, *w_rel_* = .96, *β_Q_* = 0.11, *β_gaze_* = 0.25, *A* = 163.92, *b* = 300.60, *t*_0_ = 70.83). The background colors indicate the participant’s actual choice (green = correct option, red = incorrect option). The model correctly predicted 95% of this participant’s choices (in the mean drift rate plot, the red 0’s indicate inaccurate predictions). Because the learning contexts were randomly ordered, the correct and incorrect options do not refer to the same underlying symbols on every trial. (C) Distributions of fitted parameters from the winning model in both experiments (kernel density plots).

Figure 2B shows the latent Q-values (top) and mean drift rates (bottom) extracted from the winning model after fitting a single participant’s data in Experiment 2. The middle row shows the participant’s proportional gaze data. This individual successfully learned to make reward-maximizing choices. The Q-values for the correct (maximizing) and incorrect (non-maximizing) options gradually diverged across trials, consistent with incremental RL. However, gaze behavior was more erratic: Although the correct option received more gaze on most trials, especially later in the task, there were some trials in which the lower-valued option was fixated longer. Occasionally, the gaze advantage for the incorrect option was large enough to outweigh its lower Q-value, causing the mean drift rate for the incorrect option to exceed the mean drift rate for the correct option (e.g., trial 48). When this happens, the model usually predicts an incorrect choice. The winning model successfully predicted 4 out of the 6 incorrect choices made by this participant after trial 30; in all four cases, the incorrect option received relatively more gaze. For comparison, a model without gaze effects failed to predict all six of the incorrect choices after trial 30. For another example participant, see Fig. S3.

Figure 2C shows the distributions of the parameter estimates in both experiments. The smaller learning rates, higher decision thresholds, and higher non-decision times in Experiment 1 might reflect the greater complexity of the learning environment (Figure 1A). In addition, the higher thresholds and non-decision times can explain the slower mean RTs in that experiment (see next section). The distributions of the drift scaling parameters *β_Q_* and *β_gaze_* are mostly above zero, suggesting that learned option values and proportional gaze independently influenced evidence accumulation. It is interesting that *βgaze* estimates were higher on average in Experiment 1, which might suggest that gaze has a stronger influence on choice in more complex environments. Finally, the distribution of the relative encoding parameter was bimodal in the second experiment: Although the majority of participants exhibited relative outcome encoding (context dependence), there were some who formed more absolute-like representations. See the Supplemental Material for parameter recovery simulation results. In particular, we find that *β_Q_*, *β_gaze_*, and *w_rel_* are highly recoverable (true/recovered parameter correlations between 0.87 and 0.97; Fig. S4 and S5).

### Choice-RT Patterns

Next, we examined the model’s ability to fit basic choice-RT patterns in both experiments. In early trials, participants selected the correct, reward-maximizing symbols at near-chance levels. By the end of the learning phase, choice accuracy was well above chance. Our model closely fit the learning curves in both experiments (Figures 3A and 3E; maroon lines). In contrast, a model with a linear linking function underestimated choice accuracy, especially in later trials (green lines). This result held across models with different gaze mechanisms and in both experiments: “Linear” models consistently underestimated accuracy (Fig. S6-S9).

**Figure 3:**
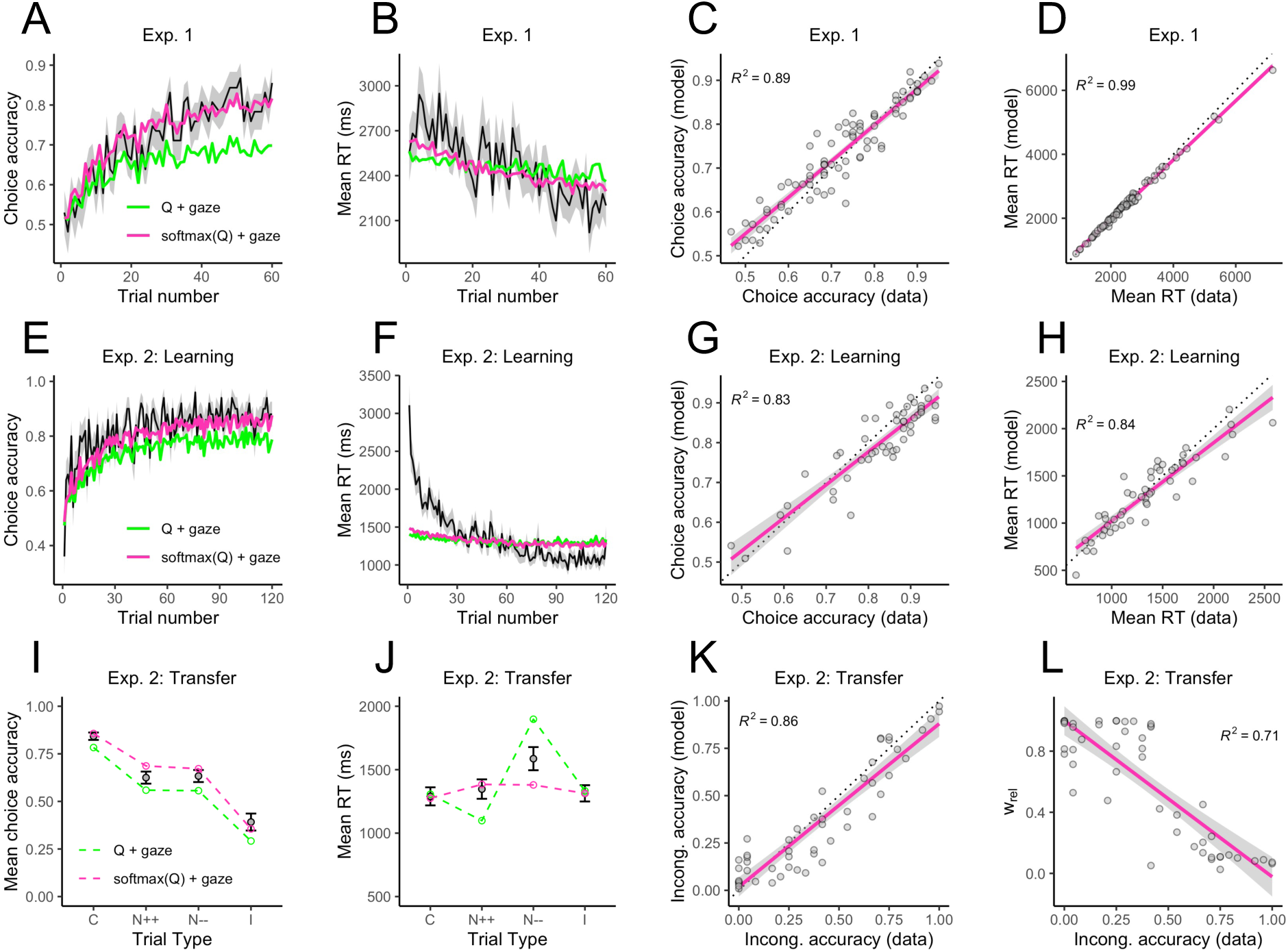
Choice-RT Effects. (A-D) Proportion of correct choices on each trial, mean RT on each trial, and relationships between empirical and fitted individual choice accuracies and mean RTs in Experiment 1. (E-H) The same as panels A-D, but for the learning phase in Experiment 2. (I-J) Mean choice accuracy and mean RT for the four types of trials in the transfer test of Experiment 2 (C = congruent, N++ = neutral with both options having high relative values, N-- = neutral with both options having low relative values, I = incongruent). (K) Relationship between empirical and fitted individual choice accuracies on incongruent trials. (L) Relationship between incongruent choice accuracies and the estimated relative encoding parameter, *w_rel_*. In all panels, error ribbons and error bars represent *±*1 standard error.

Choices also tended to become faster across the learning phase (Figures 3B and 3F). Although our model predicts this, the predicted effect was weaker than the empirical effect, especially in Experiment 2. In our model, choices become faster across trials because the mean drift rate for the correct option increases in magnitude as the underlying Q-values diverge (e.g., Figure 2B). But faster RTs could also result from individuals reducing their response caution as they grow more accustomed to the task. In RL-SSMs, this can be implemented by allowing the decision threshold to decrease across trials ^25^. Indeed, a modified version of our model with a trial-dependent, decreasing decision threshold fit the aggregate RT curve better (see Appendix B and Fig. S7). Thus, the assumption of a static decision threshold may need to be abandoned in future work if the goal is to closely fit RT curves.

Our model also accounts for heterogeneity across individuals, explaining 89% of the variance in individual choice accuracies and 99% of the variance in individual mean RTs in the learning phase of Experiment 1 (Figures 3C and 3D). The percentages were 83% and 84%, respectively, in the learning phase of Experiment 2 (Figures 3G and 3H). The only model parameter that was a significant predictor of choice accuracy in Experiment 1, controlling for the rest, was the decision threshold: Higher thresholds were associated with greater accuracy (Table S1). In Experiment 2, higher learning phase accuracy was predicted by higher learning rates, lower start point upper bounds, and higher thresholds (Table S2). Several model parameters were related to response times: Longer mean RTs were associated with lower values of *β_Q_*, lower values of *β_gaze_*, lower start point upper bounds (Experiment 1 only), and higher decision thresholds (Tables S3 and S4). Smaller values of *β_Q_* and *β_gaze_* lead to smaller drift rates and thus slower evidence accumulation, while higher thresholds and lower start point upper bounds lengthen the average distance that the accumulators must travel to reach the threshold. Taken together, slower speeds and larger distances lead to longer mean RTs.

Next we turn to the transfer test in Experiment 2, which was designed to reveal context-dependent valuation in RL. Recall that in the learning phase, each symbol was either the “better” option in its original context or the “worse” option. The transfer test involved choices between all possible pairwise combinations of symbols. Thus, the transfer choices can be categorized into four types: A higher-valued “better” option paired with a lower-valued “worse” option (Congruent trials), two “better” options (Neutral++ trials), two “worse” options (Neutral-- trials), or a higher-valued “worse” option paired with a lower-valued “better” option (Incongruent trials). Our model successfully captured the characteristic pattern of context-dependent valuation: High choice accuracy on Congruent trials, low accuracy on Incongruent trials, and intermediate accuracy on Neutral trials (Figure 3I). It provided a close fit to the mean RTs in three of the four categories; however, it underestimated the longer RTs on Neutral-- trials (Figure 3J). Interestingly, the “linear” model (green line) underestimated RTs on Neutral++ trials, where both options should have higher Q-values and drift rates, and overestimated RTs on Neutral-- trials, where both options should have lower Q-values and drift rates (that is, assuming *w_rel_* ≫ 0). The non-linear models do not exhibit this behavior because the softmax removes any distinction between two high Q-values and two low Q-values: only the difference matters. However, it should be noted that none of the models perfectly captured the pattern of mean RTs across the transfer choice categories (Fig. S12).

Lastly, there was considerable individual variability on Incongruent trials: Some participants were highly accurate in choosing the correct option, but many performed well below chance. The model explained 86% of the variability across individuals (Figure 3K). As expected, *w_rel_* was the strongest predictor of Incongruent trial accuracy out of the model parameters, with higher *w_rel_* associated with lower accuracy (Figure 3L). However, Incongruent trial accuracy was also negatively associated with *β_Q_* and the start point upper bound, and positively associated with decision thresholds (Table S5).

### Gaze Effects

Gaze effects were examined as follows. First, we computed, for each participant and each trial, the proportional amount of time that each symbol was fixated during the time period from trial onset to choice. For example, if on a particular trial there were three fixations on the left symbol with durations of 260, 136, and 180 ms, and one fixation on the right symbol with a duration of 224 ms, the proportional gaze for the left symbol would be (260 + 136 + 180)/(260 + 136 + 180 + 224) = .72 and the proportional gaze for the right symbol would be 224/(260 + 136 + 180 + 224) = .28. Non-symbol fixations and fixations on unavailable symbols (Experiment 1) were excluded from the denominator. If neither of the available symbols was fixated, both received a proportional gaze score of 0. We then computed the difference between the proportional gaze scores for the correct (higher valued) and incorrect (lower valued) symbols on each trial. This difference score ranges from 1 to -1, with positive values indicating a gaze advantage for the correct symbol, and negative values indicating a gaze advantage for the incorrect symbol.

Previous studies have shown that relative gaze predicts choice: the longer an item is fixated, the more likely it is to be chosen ^14,15,18–20^. Based on this, we should expect higher choice accuracy on learning phase trials where the correct option received relatively more gaze than the incorrect option. We grouped the gaze difference scores defined above into five quintiles, separately for each participant, and plotted the mean choice accuracy and mean RT in each quintile. Choices in the learning phase were not only more accurate, but also faster when the correct option received a greater proportion of the total fixation time, consistent with a drift rate effect (Figures 4A, 4B, 4E and 4F). Our model (maroon lines) captured these effects much better than a model without gaze bias (blue lines). In the Supplemental Material, we show that while models with multiplicative gaze mechanisms fit better than the no-gaze models, they tended to exaggerate the effects (Fig. S13-S15). Our model was also able to explain 64% (Exp. 1) and 45% (Exp. 2) of the variability across individuals in the magnitude of the gaze effect, measured as the difference in choice accuracy between the 5th gaze quintile and the 1st gaze quintile (Figures 4C and 4G). Individuals with higher values of the *βgaze* parameter tended to show larger gaze effects (Figures 4D and 4H).

**Figure 4:**
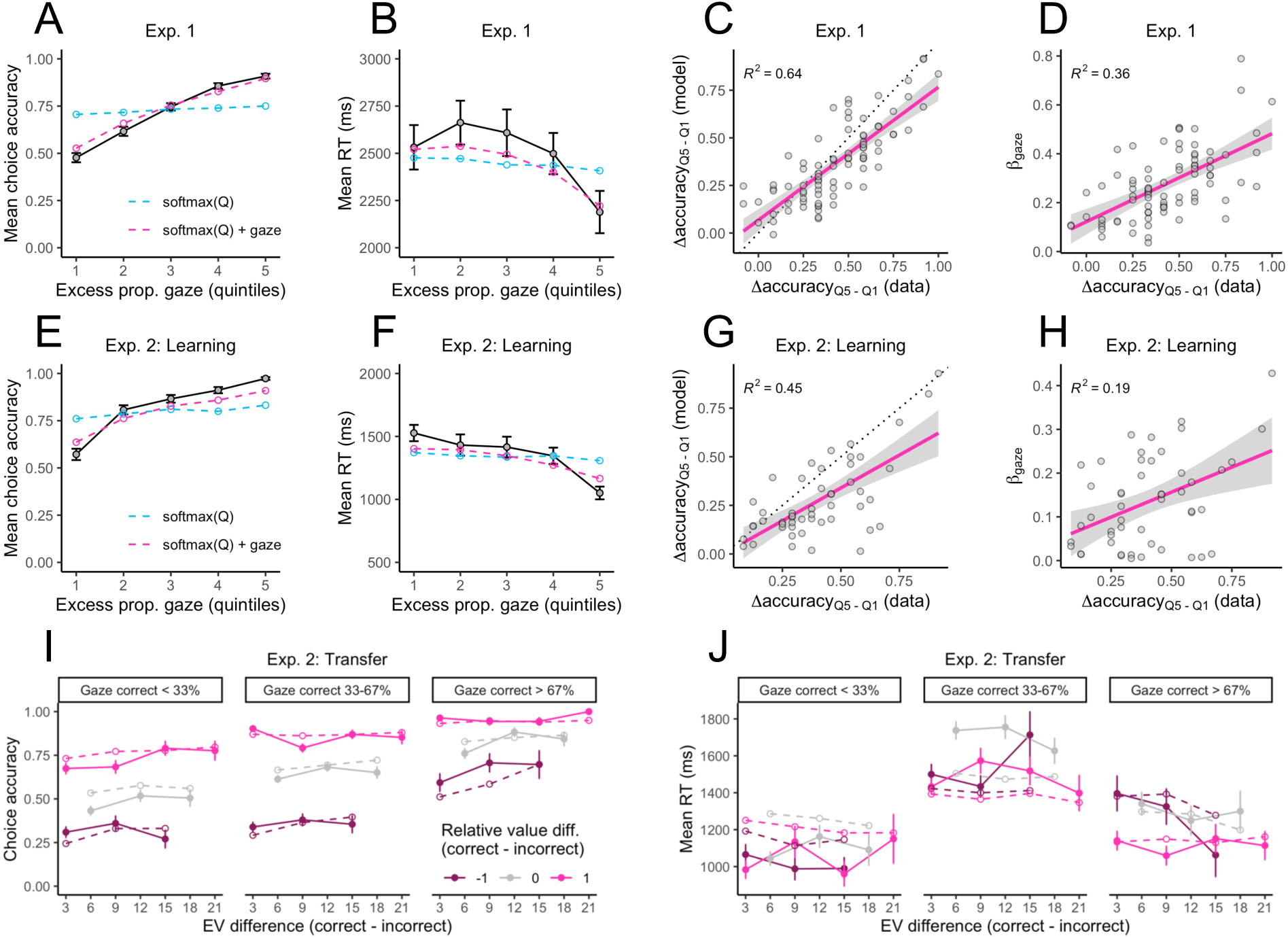
Gaze Effects. (A-D) Experiment 1 gaze effects. (A) Mean choice accuracy and (B) mean RT as a function of the excess proportional gaze on the correct option, broken up into five quintiles. The quintiles were constructed by taking the difference between the proportional gaze for the correct (higher valued) and incorrect (lower valued) symbols on each trial, sorting the difference scores, and dividing them into five equal-sized bins, separately for each participant. The higher the quintile, the longer the correct option was fixated relative to the incorrect option. (C) Relationship between empirical and fitted gaze effects at the individual level, defined as the difference in accuracy between the most extreme gaze quintiles. (D) Relationship between individual gaze effects and the gaze drift scaling parameter, *βgaze*. (E-H) Experiment 2 gaze effects in the learning phase (same as panels A-D). (I-J) Experiment 2 gaze effects in the transfer test. (I) Proportion of correct choices and (J) mean RT as a function of the expected value difference between the available options, the relative value difference, and the proportional gaze on the correct, higher-valued option. Dashed lines show model fit.

On average, the correct option had a significant proportional gaze advantage (estimated intercept = 0.059 [Exp. 1] and 0.20 [Exp. 2], ps < .001) that increased across the learning phase (trial number: slope = 0.018 [Exp. 1] and 0.058 [Exp. 2], ps < .001) and was greater for choices involving options with larger value differences (EV difference: slope = 0.035, p < .001 [Exp. 1]; linear mixed-effects regressions on proportional gaze difference scores). Thus, there is a potential confound: When the subjective values of the available options are more distinct, either because the decision maker has had more experience with the task (a learning effect) or because the underlying reward distributions are spaced far apart (a difficulty effect), the better option will not only be selected faster and with greater probability, but may also receive greater attention precisely because it is much more valuable. The important question for us is whether gaze affects choice after controlling for learning and difficulty effects.

To test this, we used logistic mixed-effects regression to model the probability of choosing the correct symbol on each learning phase trial as a function of trial number, the EV difference between the available options (correct minus incorrect; Experiment 1 only), the overall EV between the available options (correct + incorrect) and the proportional gaze difference for the available options (correct minus incorrect). The two-way interactions with trial number were also included. EV difference was not included in the model for Experiment 2 because it was constant across trials (ΔEV = 3). In both experiments, the proportional gaze difference was a significant predictor of choice accuracy even after controlling for the other predictors (gaze difference: slope = 1.076 [Exp. 1] and 1.33 [Exp. 2], ps < .001). The gaze effect weakened across trials in Experiment 1 (trial number × gaze difference: slope = -0.14, p = 0.022), but not in Experiment 2 (slope = -0.058, p = 0.41). Taken together, these results indicate that relative gaze influenced learning phase choice accuracy over and above the effects of trial number, the separation between option values, and the sum of option values.

A parallel linear mixed-effects model was performed with log-transformed response time (log-RT) as the dependent variable. In both experiments, greater gaze toward the correct option predicted faster responses even after accounting for the other predictors (gaze difference: slope = -0.031 [Exp. 1] and -0.071 [Exp. 2], ps < .001). The negative effect of gaze on log-RT strengthened across trials in Experiment 1 (trial number × gaze difference: b = -0.0097, p = 0.065), but weakened across trials in Experiment 2 (slope = 0.027, p = 0.0018). The key takeaway is that, as with choice accuracy, relative gaze influenced response times in the learning phase over and above the effects of trial number, EV differences, and overall EV.

To analyze gaze biases in the transfer test of Experiment 2, we used logistic mixed-effects regression to model the probability of choosing the correct, maximizing symbol on each trial as a function of the difference in expected values, the difference in relative values, the overall (summed) expected value, the overall relative value, and the difference in proportional gaze between the available symbols (correct minus incorrect). For this analysis, relative value was based on a symbol’s rank within its original learning context (0 = worse option, 1 = better option). The relative value difference can then take values of +1 (Congruent trials), 0 (Neutral trials), or -1 (Incongruent trials).

Relative value differences had a large impact on choice accuracy in the transfer test (slope = 1.20, p < .001), more so than expected value differences (slope = 0.27, p = .010). There was also a robust gaze effect: Holding values constant, accuracy increased with the proportional gaze allocated to the correct option (Figure 4I). In the mixed-effects model, proportional gaze difference was a significant predictor of choice accuracy over and above the other predictors (slope = 0.89, p < .001). Our computational model closely reproduced the combined effects of absolute expected values, relative values, and gaze asymmetries on transfer choice accuracy at the group level (Figure 4I).

Mean RTs in the transfer test were noisier and more difficult for our model to fit. Visual inspection of Figure 4J suggests two major trends: 1) Slower RTs on Neutral trials (relative value difference = 0) compared to Congruent and Incongruent trials (relative value difference = ±1), and 2) slower RTs on trials with a more even gaze allocation between the two symbols. Thus, we analyzed log-RTs using linear mixed-effects regression with the same predictors as the model for choice accuracy, except that the *unsigned* relative value difference and *unsigned* proportional gaze difference were used instead of the signed versions. Unsigned proportional gaze difference was a significant negative predictor of log-RT, controlling for the other predictors (slope = -0.08, p < .001). Unsigned relative value difference (slope = -0.05, p < .001) and overall relative value (slope = -0.05, p < .001) were also significant negative predictors of log-RT. Thus, RTs were slower whenever relative values or gaze proportions were (approximately) equal. Our computational model qualitatively captures these patterns. In fact, it can be easily shown that the model predicts the slowest RTs when both options have equal Q-values and equal gaze allocations. However, as previously discussed, the model struggled to predict the negative effect of overall relative value (i.e., slower RTs when both options have low relative value).

## DISCUSSION

In this study, we introduced a reinforcement learning and sequential sampling model (RL-SSM; ^1^) that can account for multiple behavioral regularities in decisions from experience. In addition to capturing choice-RT patterns and learning effects, our model uses exogenous gaze data—the proportional fixation time for each option—to constrain the evidence accumulation process on each choice trial ^14,15,19,20^. This allows it to account for gaze effects that would be missed by existing RL-SSMs ^25–29^, including the tendency for people to choose the options that they look at more ^14,15,18–21^. Although this effect has been documented in RL settings ^9,19^, this study is the first, to our knowledge, to incorporate gaze biases into a full RL-SSM that simultaneously captures learning and choice-RT effects.

Constraining our model with gaze data enables it to make better predictions. In particular, it correctly predicts that reward-maximizing options are more likely to be selected when they receive relatively more gaze prior to choice. Further, there were occasional trials in which a participant would fixate longer on the non-maximizing option before selecting it. Models that do not incorporate gaze data would misattribute these trials to random decision noise or other cognitive functions; however, our model was able to correctly predict many of these erroneous choices due to the additive effect of relative gaze on mean drift rates (see Figure 2B and Fig. S3 for examples). Our model also predicts gaze effects on choice RT: Choices were generally faster when the higher-value option received a greater share of the total fixation time. This effect is another direct consequence of the model’s additive gaze mechanism. Thus, the results of this study demonstrate the clear benefits of incorporating eye gaze data into models of experience-based decision making ^8,9^.

Central to our model is the linking function that maps learned Q-values and relative gaze onto mean drift rates. The linking function encodes two key assumptions: First, Q-values are mapped non-linearly onto mean drift rates; and second, the effect of gaze is additive and independent of learned values. Previous studies have found support for a non-linear mapping between option values and drift rates in sequential sampling models ^20,26,28,29^. The non-linearity seems to be important for capturing choice accuracy rates, as linear versions of our model consistently underestimated accuracy. The softmax mapping causes the mean drift rate for a given option to vary inversely with the learned values of the other available options. As noted by ^28^, the softmax mapping could then be interpreted as a kind of lateral inhibition between the available choice options. Lateral inhibition allows models with separate racing accumulators to emulate diffusion models, which accumulate the *relative* evidence for choosing one alternative over the other ^12^. Thus, we would expect our model to behave similarly to diffusion models despite using separate racing accumulators.

Note that the relativization of mean drift rates via the softmax is separate from the relativization of outcomes in the RL module, which is entirely controlled by the *w_rel_* parameter (Eq. 4). Relative outcome encoding allows the model to account for a specific value learning bias; that is, the tendency for learned Q-values to reflect context-dependent ranks, rather than absolute expected values ^31,41,42^. The softmax mapping, on the other hand, ensures that mean drift rates depend on the relative differences among Q-values, regardless of the degree to which the Q-values reflect absolute or relative outcomes. The former operates at the outcome encoding stage, and the latter at the decision stage. The combination of these separate relativization mechanisms allow the model to produce and fit a wide range of behavioral effects.

Our model comparison favored an additive gaze mechanism over a multiplicative gaze mechanism, consistent with some prior work ^19,21^; however, several other studies have favored a multiplicative account ^22^. Psychologically, a multiplicative gaze mechanism might suggest that directly fixating on an option facilitates the computation of its value, or the integration of past outcomes sampled from memory ^43,44^. Such a facilitation effect would benefit high-value options more than low-value options, as assumed by multiplicative gaze models ^22^. An additive gaze mechanism could instead reflect the influence of factors such as physical salience ^21,45^, estimation uncertainty ^9^, spatial location ^46^, or even random noise, all of which can bias choice independently of value. Both mechanisms qualitatively captured the major gaze effects observed in our experiments, but the multiplicative models tended to exaggerate the effects (Fig. S13-S15; see ^21^ for a similar finding). The additive gaze model provided a slightly better fit and made more accurate one-step-ahead predictions on average. However, it is important to note that additive gaze in a racing accumulator framework may not behave exactly like additive gaze in a diffusion model like the aDDM ^14^. Future research should compare additive and multiplicative gaze mechanisms across both accumulator and diffusion-based RL-SSMs using, e.g., methods found in ^37^.

Although the model with a non-linear linking function and additive gaze effect was favored on average (Figure 2A), some participants were better described by a linear linking function or by a multiplicative gaze mechanism. Additional analysis indicated that these participants exhibited different effects of value and gaze on choice and RT. For example, participants in the first experiment whose response times were faster when choosing between high-value options tended to be better described by a model with a linear relationship between Q-values and mean drift rates. Participants who exhibited a stronger effect of gaze when choosing between high-value options tended to be better described by a multiplicative gaze model (Appendix C). Both of these effects have been doc-umented in the literature ^22,47,48^; however, they were not prominent enough in our data to give models with linear linking functions or multiplicative gaze an advantage over the winning model at the group level. Future work should consider ways of combining alternative mechanisms in the same model, with parameters to arbitrate between them. For example, it may be possible to construct a linking function that interpolates between the linear and softmax mappings, so that the same model could produce and fit a wider range of behaviors.

Our model has a number of practical advantages. First, because it uses linear ballistic accumulation ^13^, it has an analytically tractable likelihood function and can be easily fit to individual choice-RT data. As there is substantial variability between individuals in the degree to which gaze influences choice ^20^, we believe the ease of fitting our model to individual data is a major benefit. Second, unlike drift diffusion models, racing accumulator models can be readily generalized to choices between three or more alternatives without any additional assumptions. Third, we showed that the model’s parameters are recoverable, even when fit to relatively small datasets (e.g., *N* = 60 observations; Fig. S4).

The present study also has important limitations. First, similar to other approaches ^14,15,19,20^, our model does not explain gaze behavior; instead, it treats gaze as an exogenous input, requiring the use of an eye-tracker to measure visual fixations. The model could be extended to also predict the relative gaze that each option receives on each trial. A good starting point would be to assume that relative gaze reflects (learned) option values ^49^—which our data support—but a more complete account would incorporate other factors such as physical salience ^21,45^ or estimation uncertainty ^9^. Second, because our model lumps fixations together in the calculation of relative gaze, it cannot account for fixation order effects, such as the tendency to choose the last-fixated option ^14^. The piecewise linear ballistic accumulator framework ^50^ may provide a solution that would enable the model to make use of the full sequence of fixations on each trial. Finally, we have evaluated our model on data from two-alternative forced choice tasks with complete feedback. We believe that increasing the number of choice alternatives and varying the nature of outcome feedback in future experiments will provide even stronger tests of the model’s core assumptions.

## MATERIALS AND METHODS

### Participants

Undergraduate students participated in two experiments in exchange for partial course credit. In the first experiment, 83 students (52 women, 29 men, 2 transgender or another gender identity; ages 18-22, M = 19.05) completed a 20-minute eye-tracking task. In the second experiment, 50 students (39 women, 11 men; ages 18-27, M = 19.1) completed a longer eye-tracking task in approximately 45 minutes. Sample sizes for both experiments were determined through power analyses designed to detect a medium-sized within-subject effect (d = 0.50) with 90% power at an alpha level of .05. Participants were informed that points earned during the learning phase (Experiment 1) or the transfer test (Experiment 2) would be converted into candy, with the conversion rate explained in the instructions. All participants were at least 18 years old, reported normal or corrected-to-normal vision, and provided informed consent. All procedures were approved by the Institutional Review Board at Binghamton University.

### Procedures

#### Experiment 1

Participants were instructed to make repeated choices between symbols with the goal of earning as many points as possible. The task was adapted from Experiment 1 in ^31^ and involved a learning phase with eight choice options grouped into two contexts of four options each. These contexts featured either a positively skewed or negatively skewed distribution of outcomes, with identical outcome ranges. Two identical options, each with an expected value (EV) of 30 points, were embedded in both contexts. The contexts were designed in this way to test competing theories of context-dependent valuation in RL (results not presented here). With four options per context, there were 12 unique option pairs across the two contexts. Each pair was presented five times, resulting in 60 total learning trials.

Context presentations were interleaved randomly across the learning phase. Symbols within a context shared a color (orange or blue) but differed in shape. The color-to-context and symbol-to-option mappings were randomized across participants. Before beginning the learning phase, participants completed five practice trials with different symbols to familiarize themselves with the setup. Each learning trial began with a drift check to align the participants gaze at the center of the screen. Then, the four symbols from one of the contexts were presented on screen, but only two symbols were available for selection (indicated by asterisks above the symbols; Figure 1A). This was done to encourage learning about all options within a context rather than simply identifying the option with the highest expected value. The four symbols, each measuring 200 × 200 pixels, were positioned 270 pixels from the screen center in one of four directions: up, down, left, or right. This corresponded to a visual angle of 8.52° between the horizontally aligned symbols and 8.58° between the vertically aligned symbols. Participants made their choice using the arrow key corresponding to the location of their preferred symbol. If a participant attempted to select one of the unavailable symbols, the trial would not advance until a valid choice was made. After a valid choice, participants returned their gaze to center for 300 ms before seeing outcomes for all four symbols (full feedback). Outcomes were overlaid on semi-transparent symbols, with the selected symbol highlighted by a white box. Feedback viewing was self-paced, and participants advanced to the next trial by pressing the space bar. Outcomes were pre-generated for each option from Gaussian distributions centered around their respective EVs with a standard deviation of 2 (rounded to the nearest integer). Although points from the chosen options were added to a running total, the cumulative score was not displayed.

At the end of the learning phase, participants were shown their total accumulated points and the maximum/minimum possible points they could have earned by choosing the correct/incorrect option on every trial. The learning phase was followed by a memory-based value estimation task (results not presented here).

#### Experiment 2

In this experiment, participants learned fixed value differences within four pairs of symbols. One symbol in each pair had a consistently higher EV than the other. Across 120 randomized learning trials, each symbol was presented 30 times within its fixed pair. On each trial, participants chose between the two symbols from a randomly selected context pair and received complete feedback, with the outcomes for both the chosen and unchosen options displayed just below each symbol. The two symbols, each measuring 300 × 300 pixels, were positioned 480 pixels to the left or right of the screen center. This corresponded to a visual angle of 15.1° between the horizontally aligned symbols. Participants were informed at the start of the learning phase to learn which symbols were most valuable. Participants made their choice between the pair by using the left or right arrow key. After each selection, participants returned their gaze to a center fixation for 300 ms before the points for both symbols were presented.

Participants then completed a transfer test that included choices between all 28 unique pairwise combinations of the eight symbols. No outcome feedback was presented during the transfer test to assess participants learned value representations. Each unique pair was presented four times for a total of 112 transfer trials. Across both phases, symbols were counterbalanced such that each appeared an equal number of times on the left and right sides of the screen. During the transfer test, participants were instructed to choose the symbol they believed would yield more points on each trial. They were also informed that the points accumulated in this phase would be converted into candy, with the specific conversion rate displayed to indicate how many candies they could earn based on their accuracy in the test. Each transfer trial began with a drift check fixation, followed by two symbols presented side by side. Using the same arrow keys as in the learning phase, participants made their selection, and the chosen symbol remained highlighted for 0.5 seconds by a white box before the next trial began.

### Eye Tracking

Fixation data was collected with SR Research EyeLink software (version 5.15). Gaze position was recorded from the right eye using a table-mounted EyeLink 1000 Plus eye tracker with the sampling rate set to 500 Hz. Stimuli were presented on a 24-inch HD LCD monitor with a 1920 × 1080 pixel resolution against a constant gray background. Participants sat approximately 100 cm from the screen with their heads stabilized in a chinrest under consistent, slightly dimmed lighting. In Experiment 1, a 9-point calibration and validation procedure was performed once at the beginning of the session. In Experiment 2, calibration and validation were performed at the beginning of the experiment and again before the start of the transfer test. Fixations were automatically identified using the EyeLink software, and rectangular areas of interest (AOIs) were defined around each symbol that measured 270 × 270 pixels in Experiment 1 and 340 × 340 pixels in Experiment 2. The average validation error was approximately 0.40° of visual angle in both experiments.

### Statistical Analysis

We analyzed gaze effects using a series of mixed-effects regression models from the “afex” package in R ^51^. Choice accuracy was analyzed with mixed-effects logistic regression, and proportional gaze difference and log-transformed RTs were analyzed with linear mixed-effects regression. All predictors were z-scored within-subject to facilitate interpretability and model convergence. We utilized a maximal random effects structure by including random intercepts and slopes for each within-subject predictor to account for heterogeneity across participants ^52^. Almost all models successfully converged under this structure. However, one model did not converge, so we removed the random effect correlations. The conclusions were not affected by whether the random effect correlations were included. Estimated coefficients and statistical tests for all mixed-effects models are provided in the Supplemental Material (Tables S7-S13).

### Computational Modeling

RL-SSMs were fit to individual choice-RT data using maximum a posteriori (MAP) estimation ^53^. MAP estimation is very similar to maximum likelihood estimation, except it uses priors to regularize the model parameters. We chose independent, moderately informative priors based on past research (e.g., realistic values for non-decision time; ^54^) and simulations of the model across its parameter space: *α* ∼ Beta(1.3, 3.7), *w_rel_* ∼ Beta(1.1, 1.1), *β_Q_* ∼ Gamma(2, 0.5), *β_gaze_* ∼ Gamma(2, 0.5), *A* ∼ Gamma(6, 100), *b_sep_* ∼ Gamma(6, 100), *t*_0_ ∼ Gamma(6, 30) (note: gamma distributions have the shape-scale parameterization). Assigning a non-negative prior to the threshold separation parameter *bsep* ensures that the start point upper bound (*A*) will not exceed the decision threshold (*b*), since *b* = *A*+*bsep*. For all fits, we fixed the drift rate standard deviation *s* to 0.1 and initialized Q-values to 0.5.

After assigning priors to the parameters, the MAP objective function becomes the sum of the log-likelihood and the log-prior (i.e., the log-posterior). The log-likelihood function for our RL-SSM is the same as that for the regular LBA model (see ^13^), but with mean drift rates computed using the linking function in Eq. 2. Log-likelihoods were summed across trials, skipping trials with RTs faster than 250 ms or slower than 10 s. We searched for parameters that optimized the MAP objective function using differential evolution optimization (NP = 100, itermax = 1000, steptol = 250) ^55^.

When simulating the RL-SSM, we used the current Q-values and proportional gaze scores for the available options to compute the mean drift rates *v_i,t_* on each trial according to Eq. 2. Start points *z_i,t_* for each accumulator were drawn independently from a Uniform(0*, A*) distribution, and drift rates *d_i,t_* were drawn independently from Normal(*v_i,t_, s*) distributions (resetting negative drift rates to zero). The time taken for the *i*th accumulator to reach the threshold can then be computed as

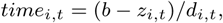

i.e., time = distance / speed. The simulated choice is given by the index *i* with the minimum *time_i,t_*, and the simulated RT is given by *t*_0_ +min*_i_*(*time_i,t_*). This procedure was repeated until the simulated RT was at least 250 ms but no longer than 10 s. In Figures 3 and 4, the model predictions were averaged across 100 simulations using each participant’s best-fitting parameters.

## ACKNOWLEDGMENTS

The authors would like to thank Andrew Dolinsky, Rishi Heggawadi, Nadiah Layne, Trevor Rosenthal, Joelle Sacks, and Elaine Yu for their assistance with data collection.

## AUTHOR CONTRIBUTIONS

W.M.H. and M.J.T. designed and performed the research; W.M.H. conducted the computational modeling analyses; W.M.H. and M.J.T. conducted the statistical analyses; W.M.H. and M.J.T. wrote the paper.

## DATA AND CODE AVAILABILITY

Data and code are available at https://github.com/william-hayes/RL-LBA-gaze-model.

## APPENDIX A

Here we describe the seven RL-SSMs that were tested. The only difference between the models is the linking function that maps learned Q-values and proportional gaze onto the mean drift rates, *v_i,t_*. Note that the additive gaze models contain one additional parameter, *βgaze*, compared to the other models. Following ^28^, we set *θ* = 50 in the softmax models.

### 1. Q Model

The simplest model that we tested maps Q-values linearly onto mean drift rates and assumes no effect of gaze:

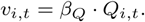

### 2. Q*gaze Model

This model also assumes a linear relationship between Q-values and mean drift rates, but with a multiplicative effect of gaze. Gaze will have a larger effect on drift rates for options with larger Q-values:

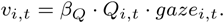

### 3. Q + gaze Model

This model assumes a linear relationship between Q-values and mean drift rates with an additive effect of gaze. The effects of Q-values and gaze, represented by *β_Q_* and *β_gaze_*, are independent:

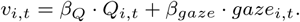

### 4. softmax(Q) Model

In this model, Q-values are non-linearly mapped to mean drift rates using the softmax function. However, there are no gaze effects:

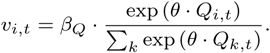

### 5. softmax(Q*gaze) Model

This model assumes a multiplicative gaze effect that operates prior to the softmax transformation (i.e., a “pre-multiplicative” effect). Gaze modulates the cached Q-values directly, and the softmax function normalizes the gaze-modulated Q-values to the unit interval:

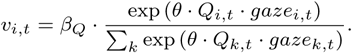

In this model, if neither option is fixated on a particular trial, such that *gaze_i,t_* = 0 for all *i*, the mean drift rate for every option will be *v_i,t_* = *β_Q_/K*, where *K* denotes the number of available options on that trial. *Importantly, this model implies that the softmax normalization must occur at the decision stage*. This is because gaze modulates the inputs to the softmax, and gaze is only measured during the decision stage.

### 6. softmax(Q)*gaze Model

This model assumes a multiplicative gaze effect that operates after the softmax transformation (i.e., a “post-multiplicative” effect). Gaze modulates the softmax-transformed Q-values:

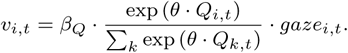

In this model, if neither option is fixated on a particular trial, such that *gaze_i,t_*= 0 for all *i*, the mean drift rate for every option will be *v_i,t_* = 0, as in the Q*gaze model.

### 7. softmax(Q) + gaze Model

The final model is the one presented in the main text. It assumes a non-linear mapping of Q-values to mean drift rates via the softmax, along with an independent, additive effect of gaze:

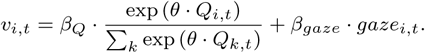

## APPENDIX B

Our primary models assume a static decision threshold; that is, that the decision maker requires the same amount of evidence to make a decision on all trials. An alternative possibility is that the decision threshold changes across trials ^25^. For example, if decision makers gradually “let their guard down,” becoming less cautious over time as they become more familiar with the task, then they might require more evidence to make a decision in earlier trials (i.e., higher threshold) and less evidence in later trials (i.e., lower threshold). The expected result would be a decrease in response times (RT) across trials. In Experiment 2, we tested a second set of models with trial-dependent (decreasing) decision thresholds to determine whether they could better capture the aggregate RT patterns. In these models the decision threshold changes across trials according to the following functional form:

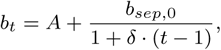

where *b_t_* is the decision threshold on trial *t*, *A* is the upper bound of the start point distribution, *b_sep,_*_0_ is the baseline (initial) decision threshold, and *δ* is a decay parameter. Note that if *δ* = 0, the model reduces to a static threshold model: *b_t_* = *A* + *b_sep,_*_0_ for all *t*. When *δ >* 0, the threshold will decrease across trials toward the start point upper bound. Higher values of the decay parameter lead to steeper decreases in the decision threshold across trials. When fitting this model to individual choice-RT data in Experiment 2, we used the following priors: *b_sep,_*_0_ ∼ Gamma(6, 100) and *δ* ∼ Gamma(1.01, 0.1) (shape-scale parameterization). As shown in Fig. S5, the models with trial-dependent decision thresholds fit the aggregate RT curves better than the models with static thresholds. The mean parameter estimates were *b_sep,_*_0_ = 313.68 (SD = 168.60) and *δ* = 0.07 (SD = 0.12), and *A* = 302.37 (SD = 142.30).

## APPENDIX C

Although the “softmax(Q) + gaze” model (Model 7) performed best on average (Figure 2A), some participants were better described by one of the other models. To shed light on these individual differences, we used the estimated individual-level coefficients from mixed effects models, which represent person-specific effects of value and gaze on choice accuracy and response time, to predict Model 7’s relative advantage for individual participants. The dependent variable in this analysis is the difference in accumulative one-step-ahead prediction error (ΔAPE) between a particular model and Model 7, where positive numbers indicate an advantage for Model 7, and negative numbers indicate an advantage for the other model. For simplicity, we only analyzed the data from Experiment 1 to avoid having to distinguish between absolute and relative values in the analysis.

First, given that the “Q + gaze” model (Model 3) predicts a negative effect of overall value on RT, while Model 7 predicts a negative effect of the value difference (Fig. S1, right panels), we expected that the individual-level coefficients for these two effects (derived from the model in Table S10) would be predictive of Model 7’s advantage over Model 3, but in opposite directions. Confirming our expectations, the slope for the effect of EV difference was negative (-84.07) and the slope for the effect of overall EV was positive (43.47) with both ps < .001 (multiple regression; adjusted *R*^2^ = .20); thus, Model 7 was better at accounting for participants who exhibited a more negative effect of EV difference and a less negative effect of summed EV on log RT.

Second, given that the “Q * gaze” model (Model 2) predicts a stronger effect of gaze on choice for options with higher overall value, while Model 7 predicts a constant gaze effect (Fig. S2), we expected that individual-level coefficients for the interaction between proportional gaze difference and overall EV would be negatively associated with the advantage of Model 7 over Model 2. We first fit a generalized linear mixed-effects model predicting the probability of choosing the correct symbol on each learning phase trial as a function of the overall (summed) EV of the available options, the proportional gaze difference for the available options (correct minus incorrect), and the interaction. Although the fixed group-level interaction was not significant (p = .97), the correlation between the individual-level interaction effects and the advantage of Model 7 over Model 2 was negative and significant (*r* = −0.35*, p* = .001). Thus, Model 7 did not perform as well for participants in the first experiment who exhibited a stronger gaze effect for higher-value choice pairs.

---

**Figure S1:**
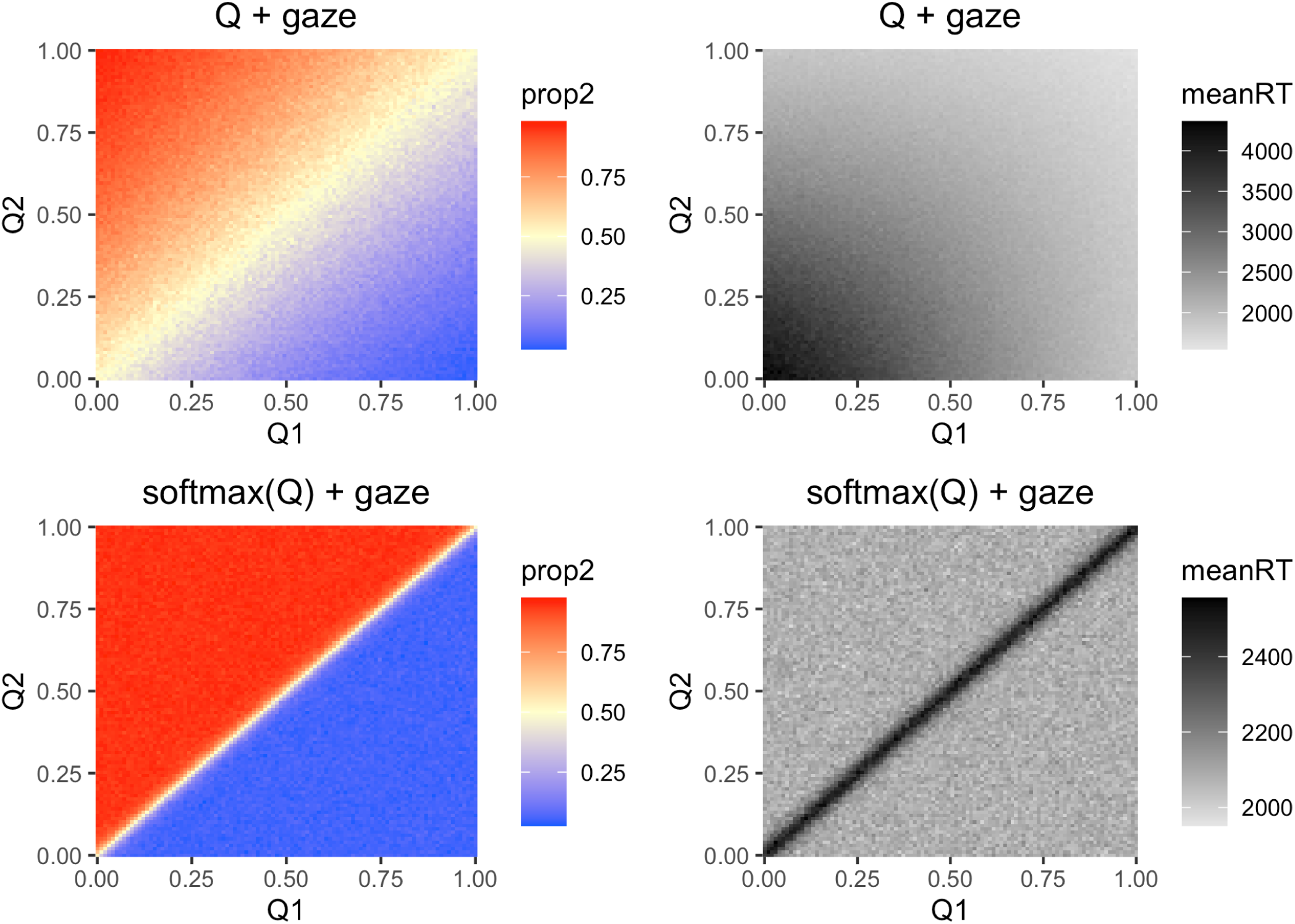
Simulated choice-RT patterns in a two-alternative choice task. Each model was simulated 500 times for each combination of Q-values. The left panels show the proportion of times the second alternative was chosen (prop2) and the right panels show the mean RT (ms) across the 500 runs for each Q-value combination. Models were simulated using the mean parameter estimates from Experiment 1 (Q + gaze: *β_Q_* = 0.32, *β_gaze_* = 0.27, *A* = 631.09, *b* = 1086.19, *t*_0_ = 133.44; softmax(Q) + gaze: *β_Q_* = 0.27, *β_gaze_* = 0.28, *A* = 613.52, *b* = 1065.47, *t*_0_ = 135.98). For both models, proportional gaze was set to 0.5 for both alternatives. The Q + gaze model exhibits a noisier decision boundary and produces mean RTs that depend on the magnitude of the Q-values, with faster RTs for larger magnitudes. The softmax(Q) + gaze model exhibits a sharper decision boundary and mean RTs that depend only on the difference between Q values, with faster RTs for larger differences.

**Figure S2:**
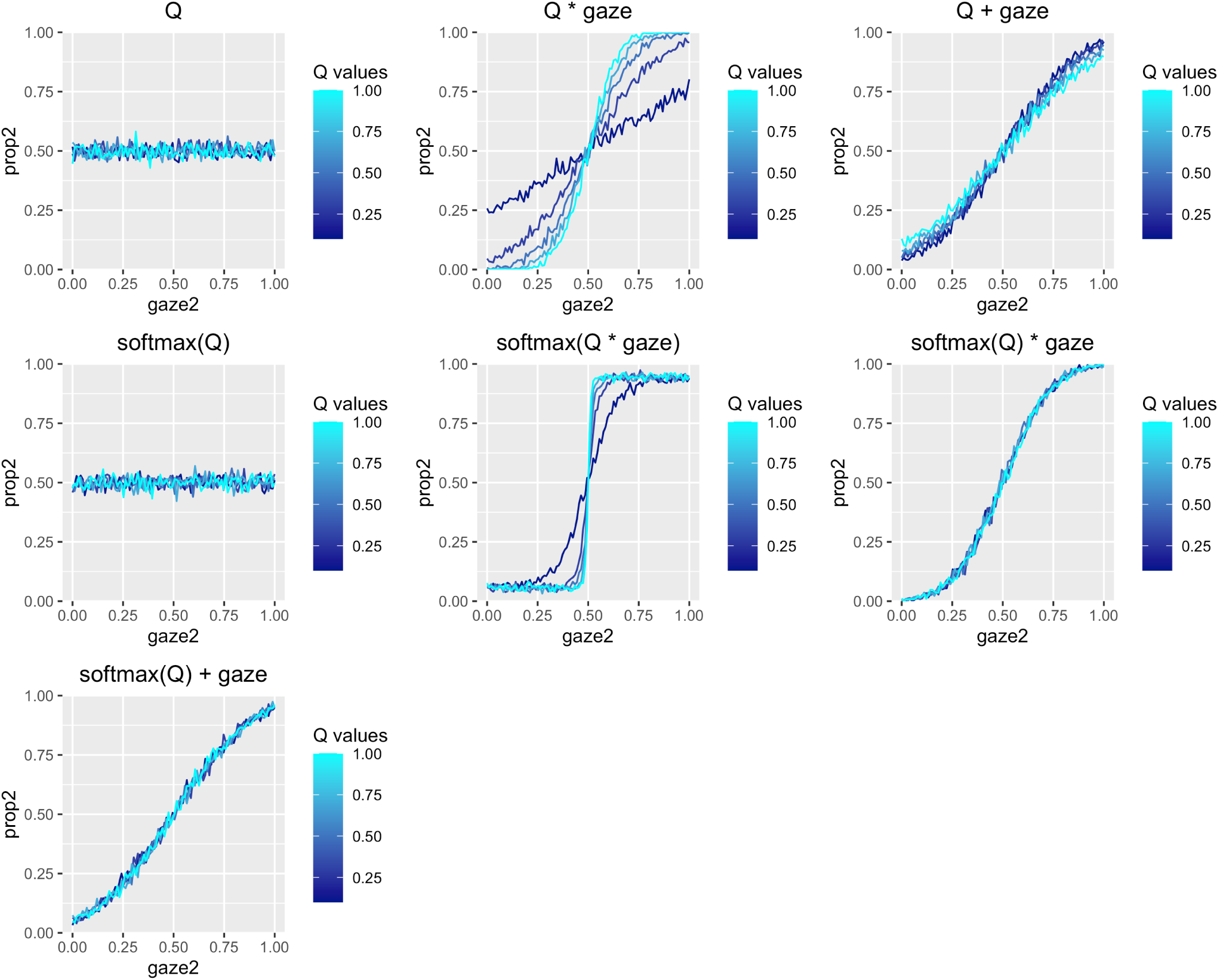
Simulated gaze effects in a two-alternative choice task. Models were simulated 500 times for each combination of Q-value magnitude (5 levels: 0.1, 0.3, 0.5, 0.7, 0.9) and proportional gaze allocated to the second alternative (100 evenly-spaced values between 0 and 1). Both options were given the same Q-value in the simulations (one of the five magnitudes listed above). Models were simulated using the mean parameter estimates from Experiment 1. The y-axis in each panel is the proportion of times the second alternative was chosen (prop2). The models without gaze data show no gaze effects. When both options have the same Q-value, the Q * gaze and softmax(Q * gaze) models exhibit stronger gaze effects when the magnitude of the Q-values is larger.

**Figure S3:**
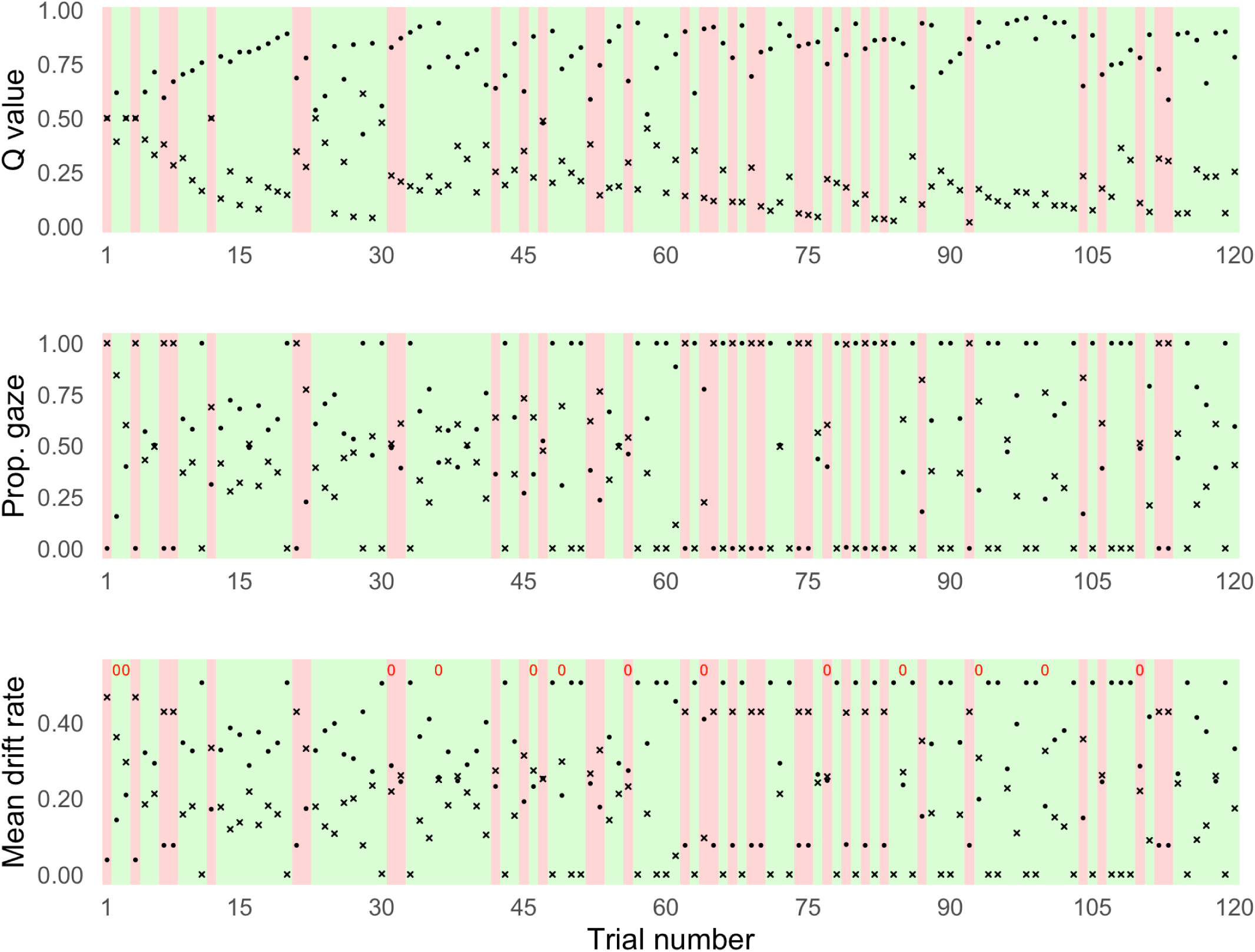
Trial-to-trial Q-values, proportional gaze data, and mean drift rates for participant 43 in Experiment 2 (parameter estimates: *α* = .25, *w_rel_* = .83, *β_Q_* = 0.08, *β_gaze_* = 0.43, *A* = 269.69, *b* = 362.98, *t*_0_ = 95.76). The background colors indicate the participant’s actual choice (green = correct option, red = incorrect option). The model correctly predicted 89% of this participant’s choices (in the mean drift rate plot, the red 0’s indicate inaccurate predictions). Because the learning contexts were randomly ordered, the correct and incorrect options do not refer to the same underlying symbols on every trial.

**Figure S4:**
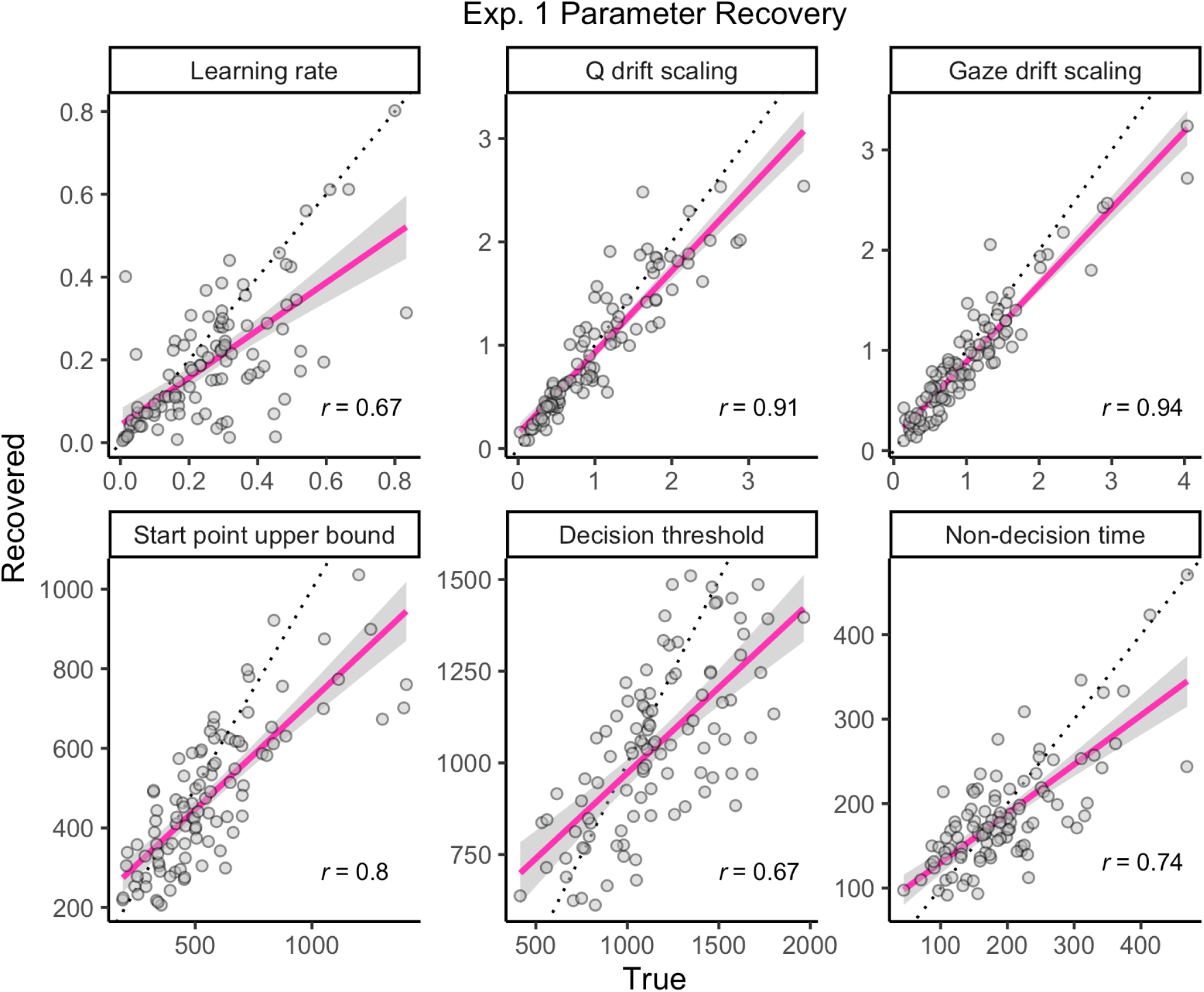
Parameter recovery in Experiment 1: The softmax(Q) + gaze model was simulated 100 times in the task from the first experiment (60 trials) using parameter values drawn from the prior distributions (see main text, *Materials and Methods*). Then, the model was fit to the 100 simulated data sets to assess its ability to recover the true, data-generating parameters. Each panel shows the relationship between the generating and recovered parameters with regression lines overlaid.

**Figure S5:**
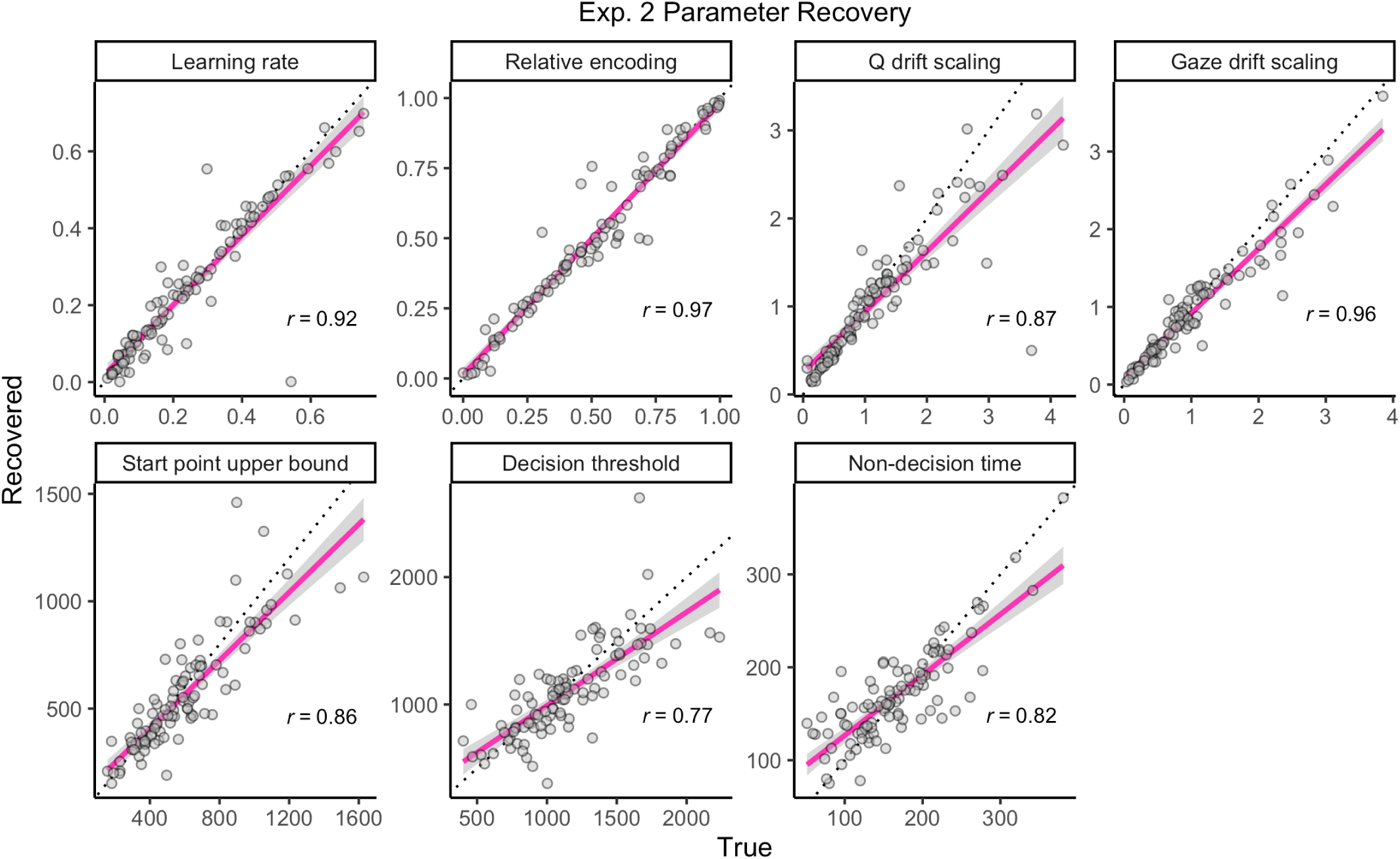
Parameter recovery in Experiment 2: The softmax(Q) + gaze model was simulated 100 times in the task from the second experiment (232 trials) using parameter values drawn from the prior distributions (see main text, *Materials and Methods*). Then, the model was fit to the 100 simulated data sets to assess its ability to recover the true, data-generating parameters. Each panel shows the relationship between the generating and recovered parameters with regression lines overlaid.

**Figure S6:**
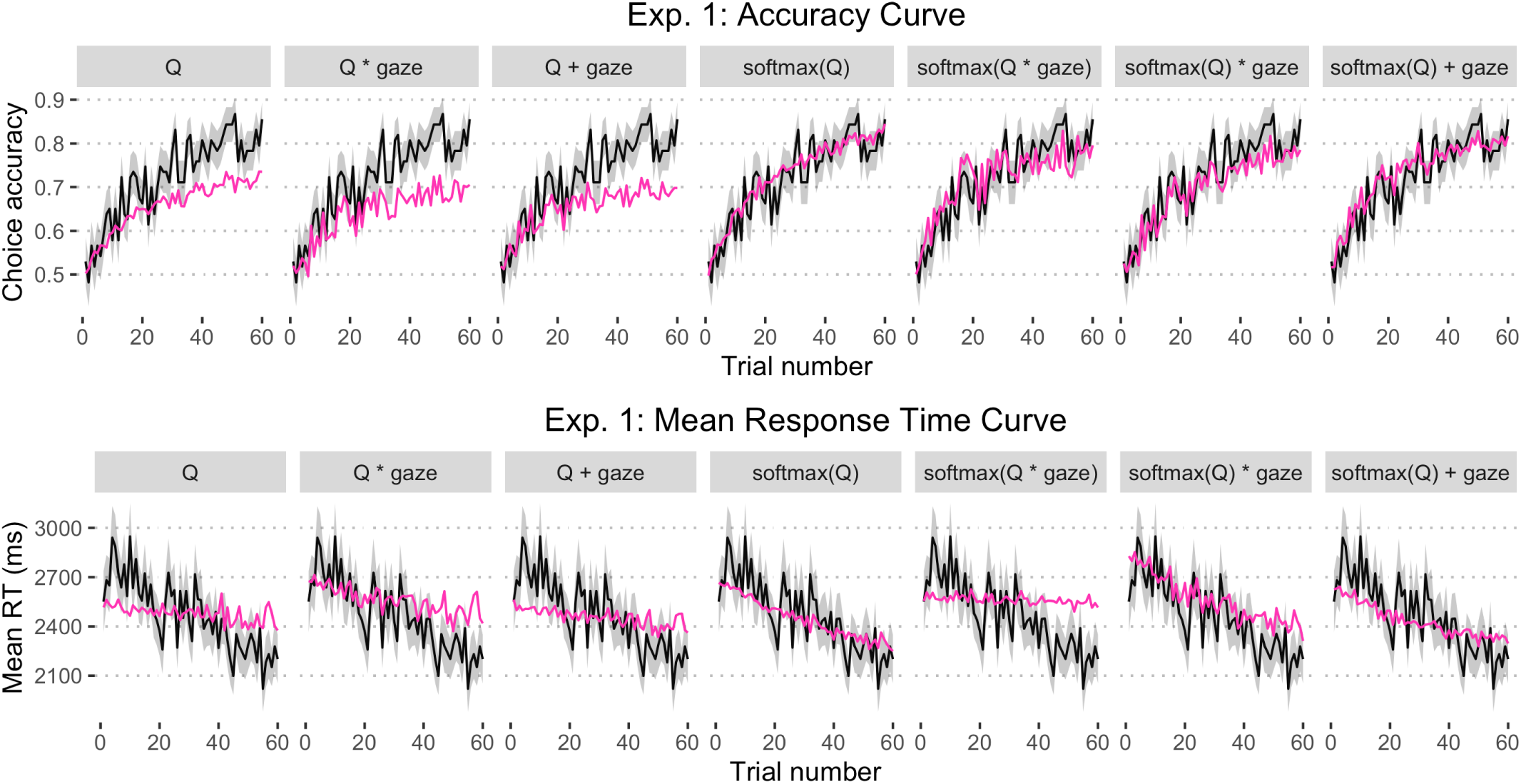
Model fit in Experiment 1: Proportion of correct choices and mean RT across learning trials. Error ribbons represent *±*1 standard error.

**Figure S7:**
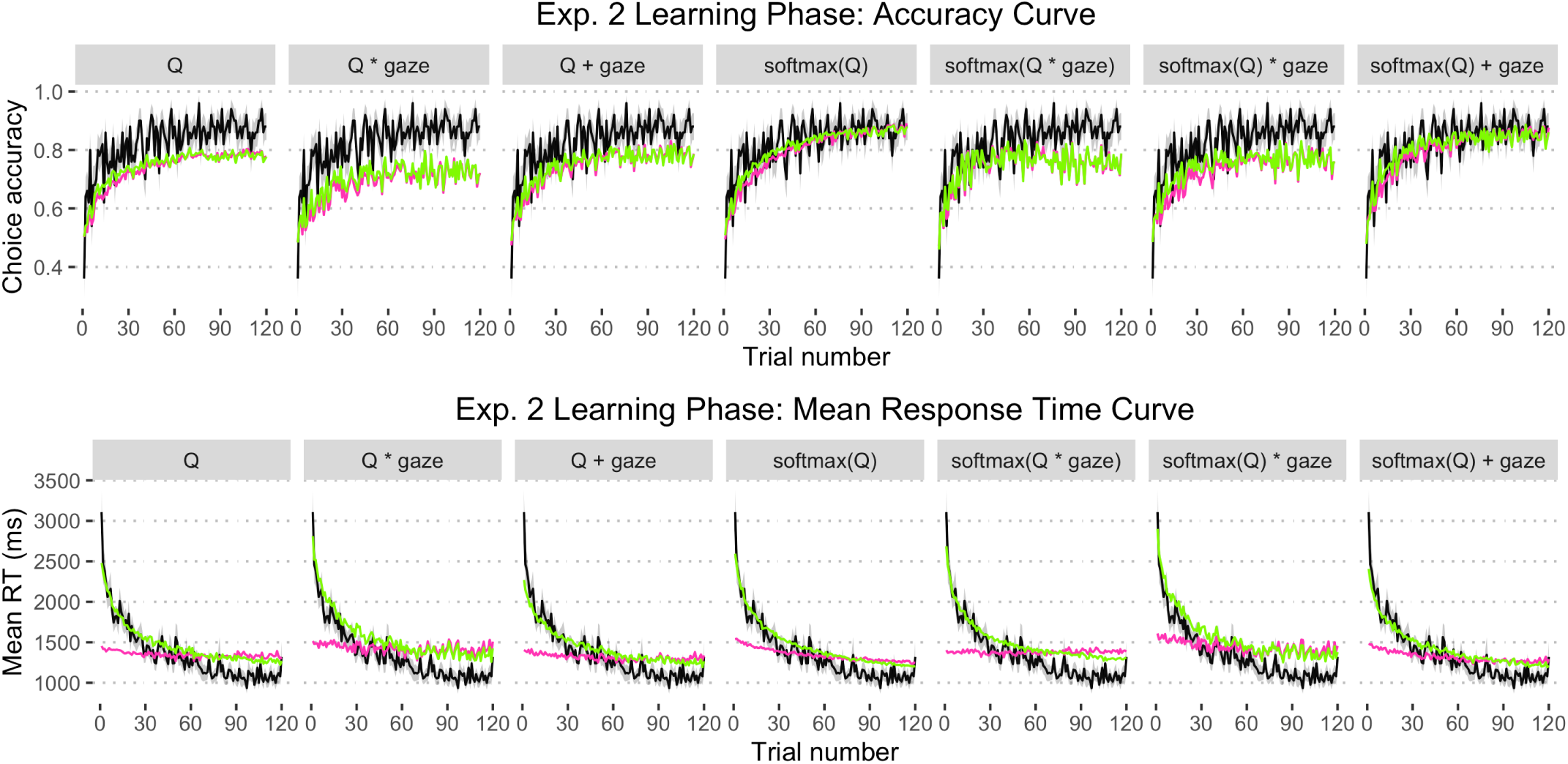
Model fit in Experiment 2: Proportion of correct choices and mean RT across learning trials. Error ribbons represent *±*1 standard error. The maroon lines show the fit of the original model with a static decision threshold. The chartreuse lines show the fit of the modified model with a trial-dependent (decreasing) decision threshold, which was better at capturing the steep, nonlinear decrease in mean RT across the learning phase.

**Figure S8:**
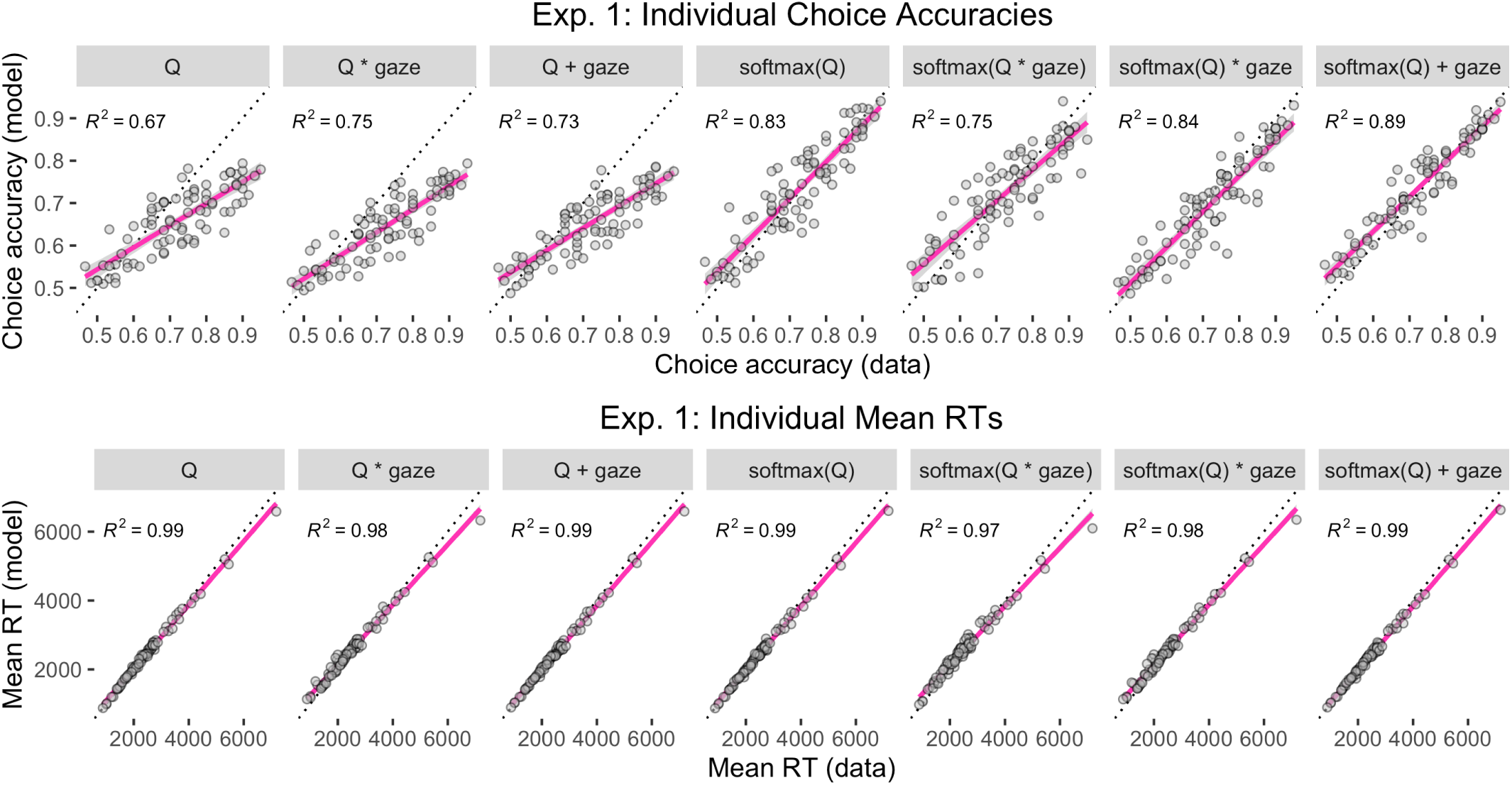
Model fit in Experiment 1: Individual choice accuracies and mean RTs.

**Figure S9:**
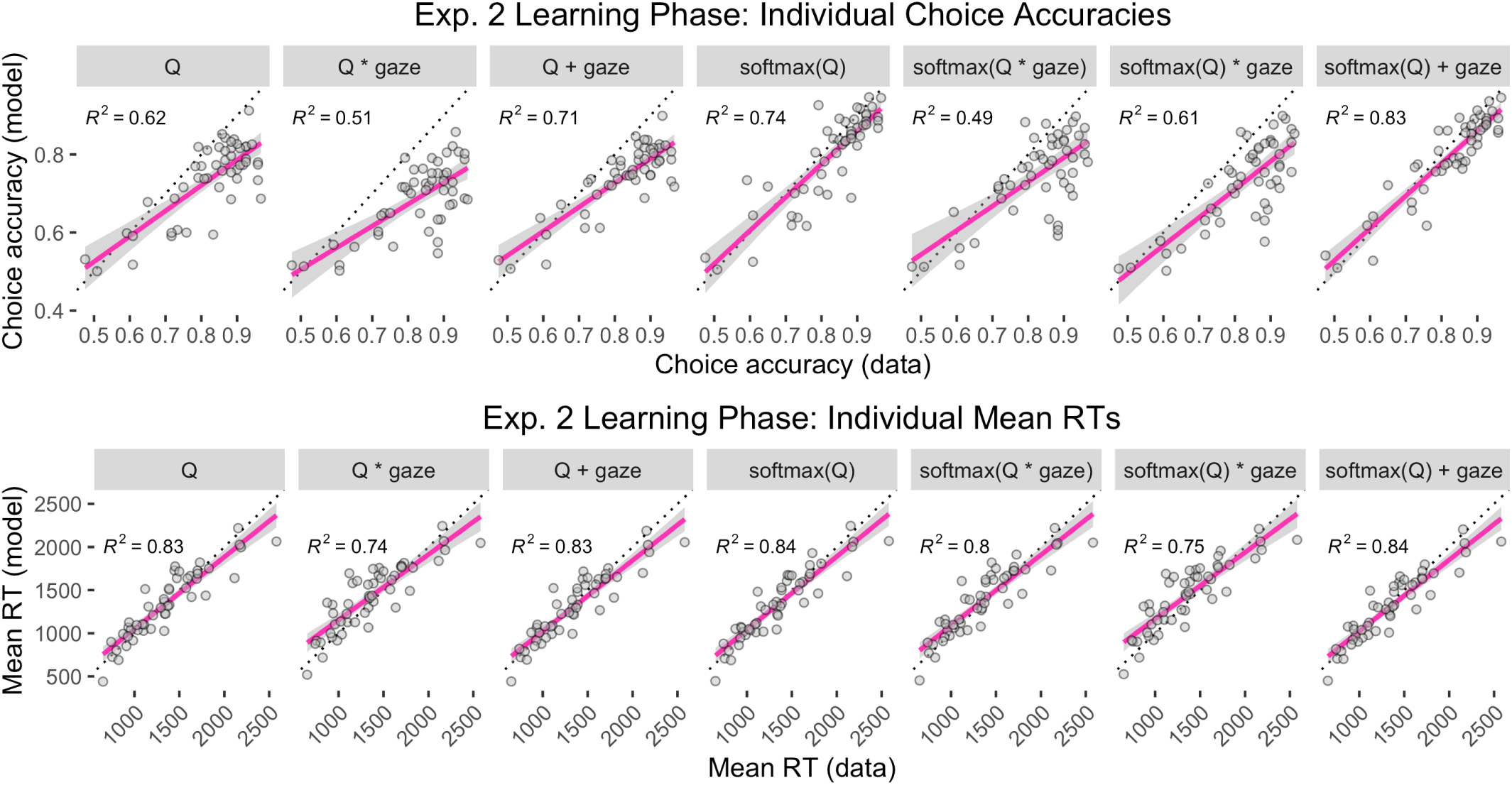
Model fit in Experiment 2: Individual choice accuracies and mean RTs in the learning phase.

**Figure S10:**
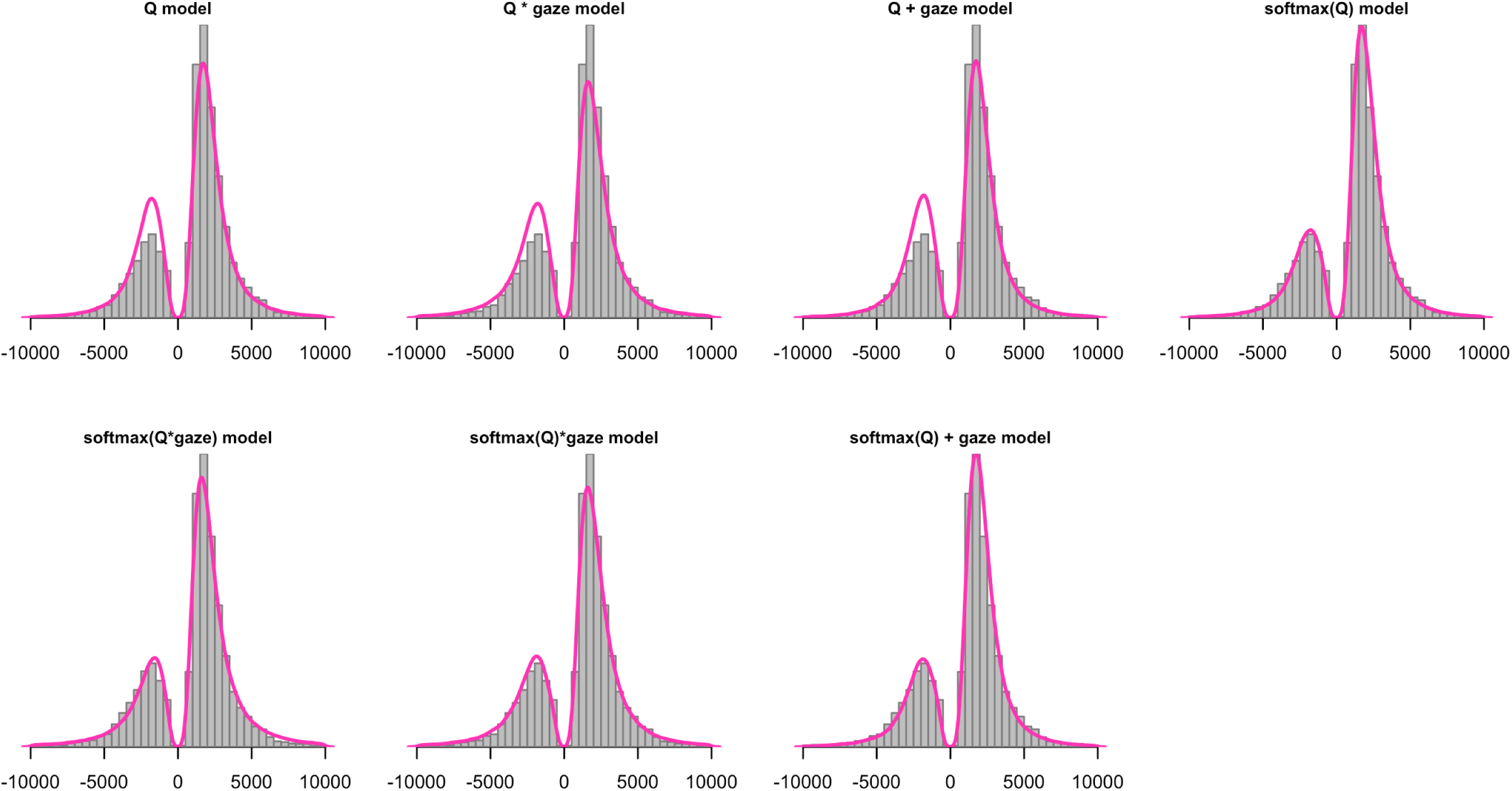
Model fit in Experiment 1: Response time distributions for correct (maximizing) and incorrect (non-maximizing) choices, pooled across participants. Incorrect RTs are negative for visualization purposes.

**Figure S11:**
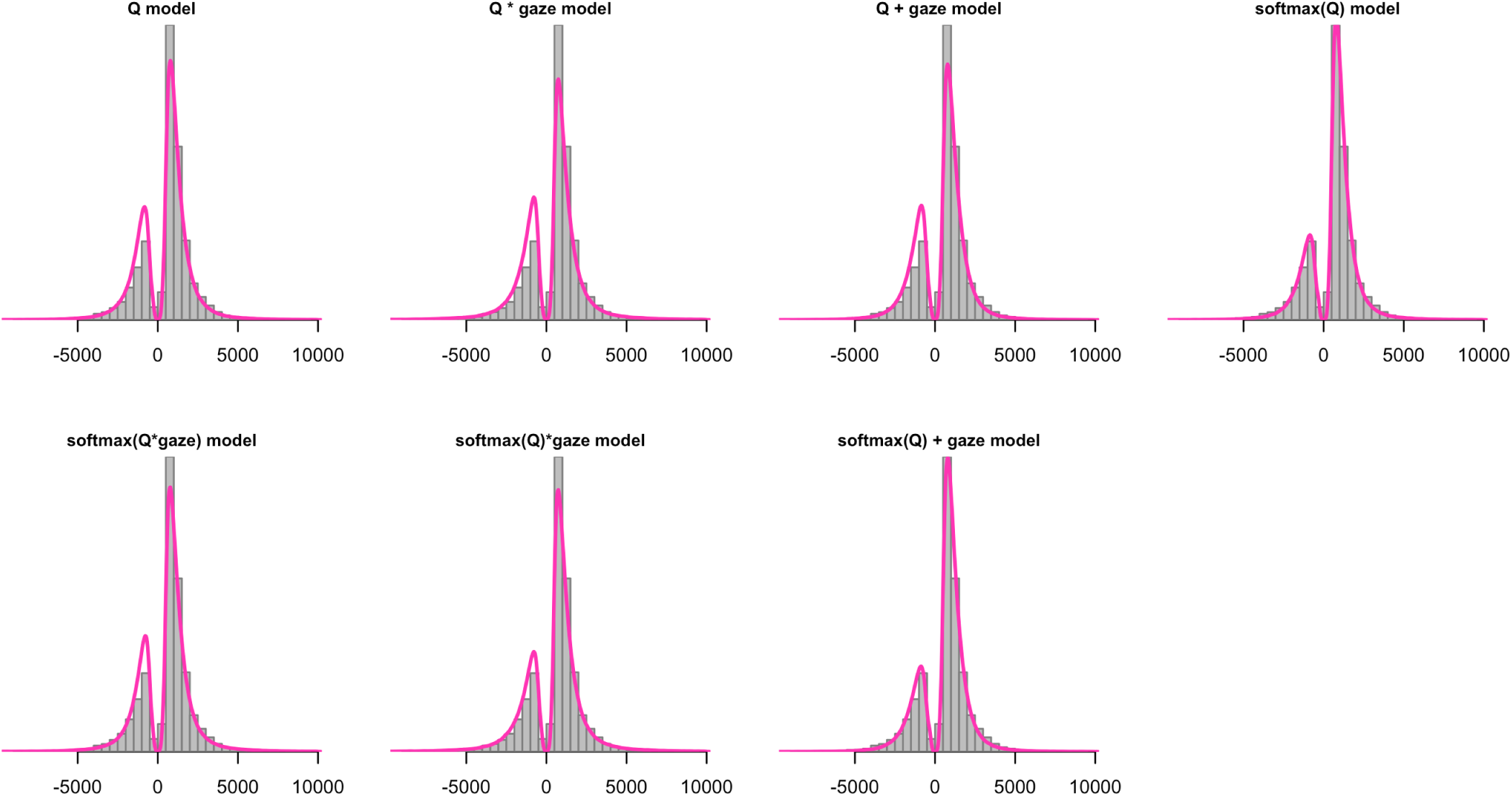
Model fit in Experiment 2: Response time distributions for correct (maximizing) and incorrect (non-maximizing) learning phase choices, pooled across participants. Incorrect RTs are negative for visualization purposes.

**Figure S12:**
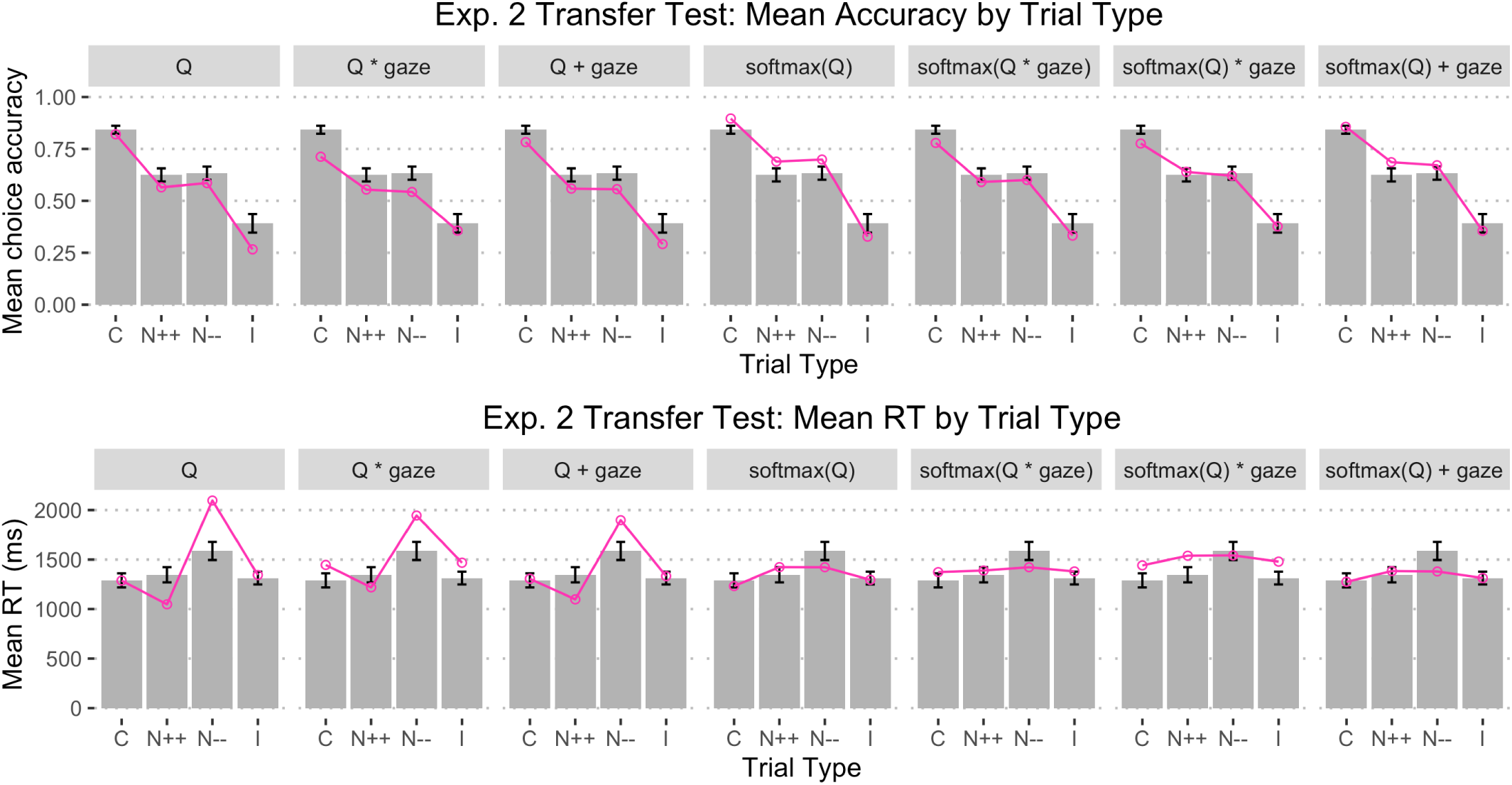
Model fit in Experiment 2: Mean choice accuracy and mean RT as a function of trial type in the transfer test (C = congruent, N++ = neutral with both options having high relative values, N-- = neutral with both options having low relative values, I = incongruent). Error bars represent *±*1 standard error.

**Figure S13:**
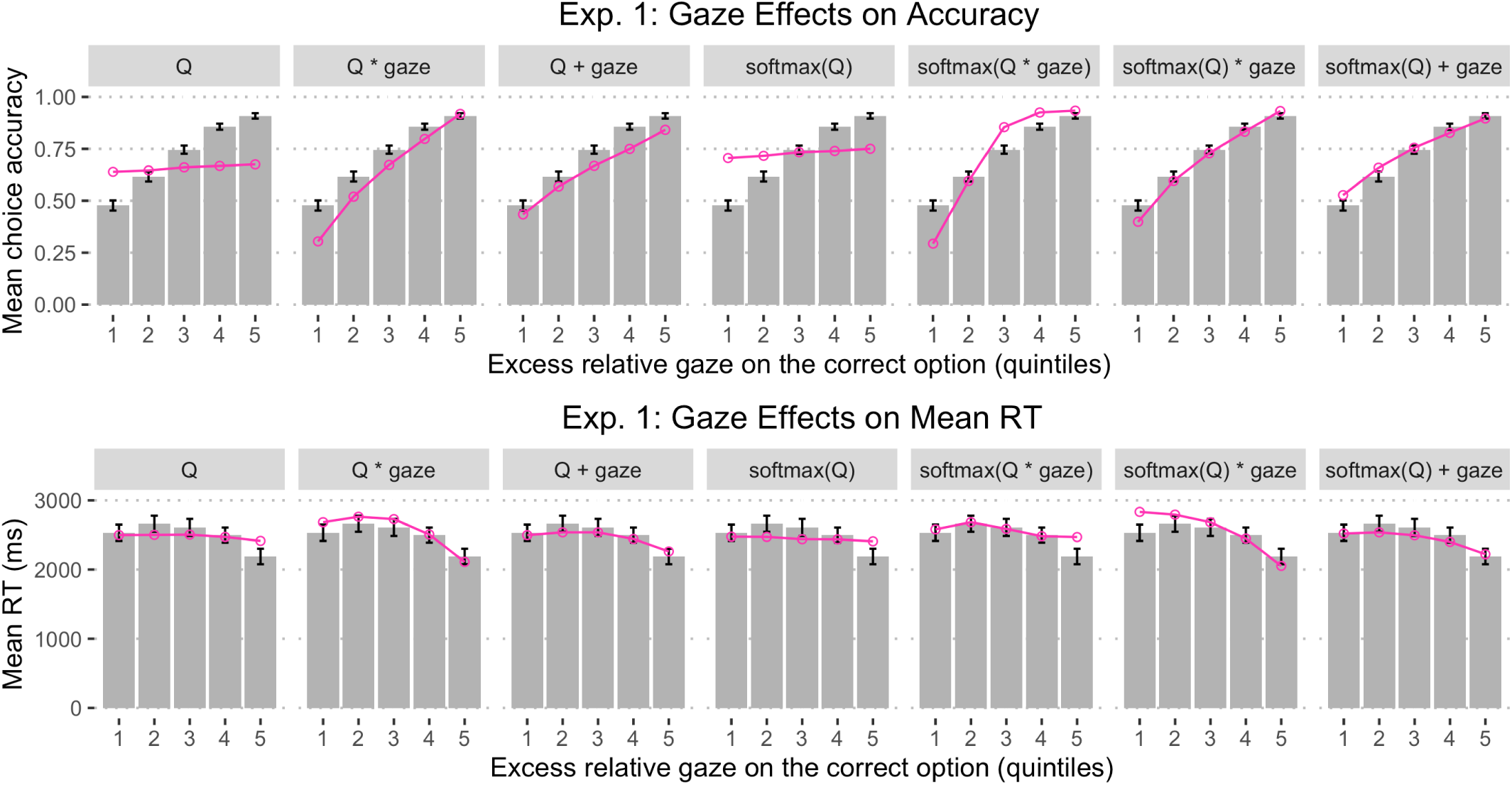
Model fit in Experiment 1: Mean choice accuracy (top) and mean RT (bottom) as a function of the excess proportional gaze on the correct option, broken up into five quintiles. The quintiles were constructed by taking the difference between the proportional gaze for the correct (higher valued) and incorrect (lower valued) symbols on each trial, sorting the difference scores, and dividing them into five equal-sized bins, separately for each participant. The higher the quintile, the longer the correct option was fixated relative to the incorrect option. Error bars represent *±*1 standard error.

**Figure S14:**
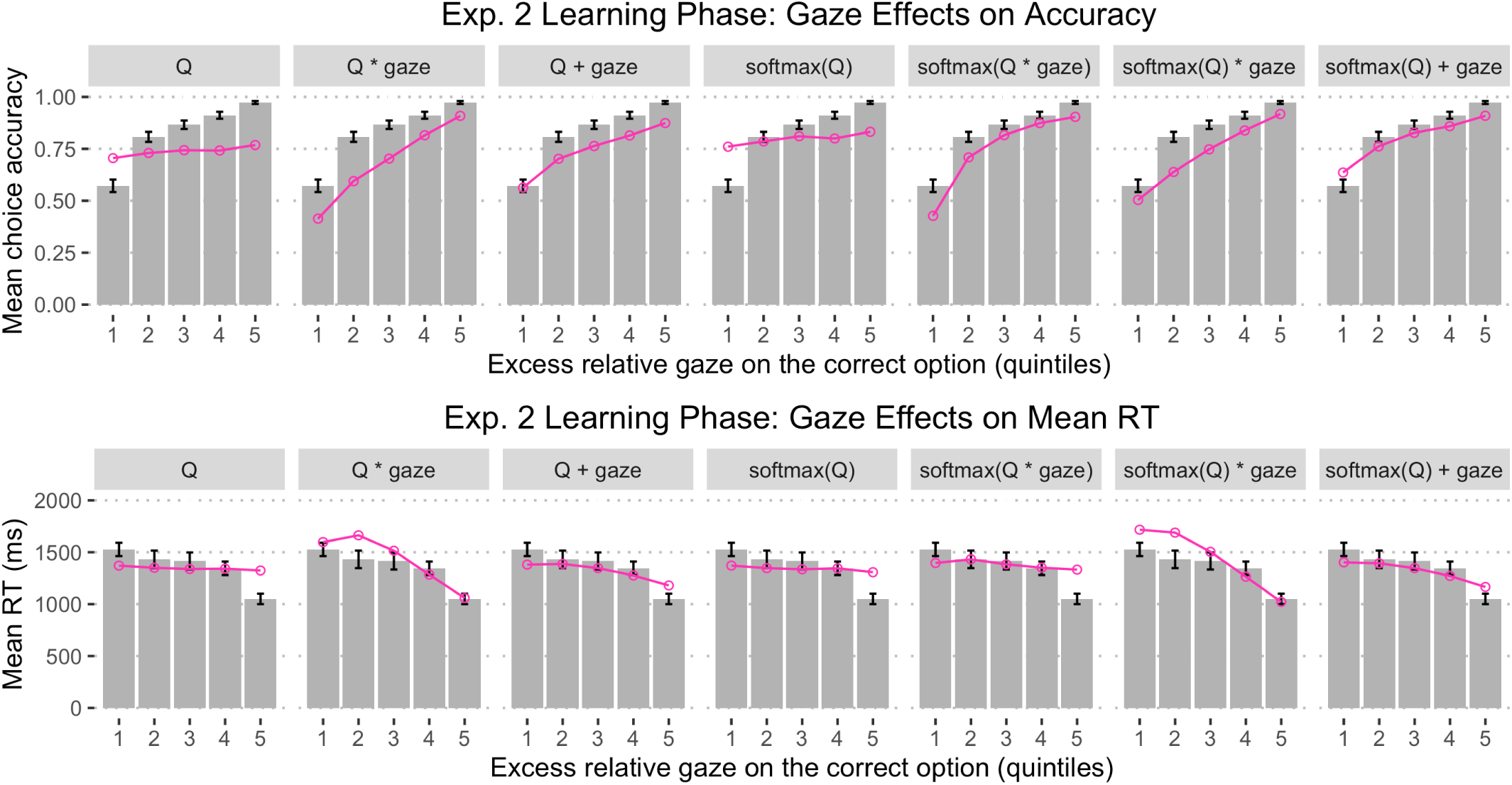
Model fit in Experiment 2: Mean choice accuracy (top) and mean RT (bottom) in the learning phase as a function of the excess proportional gaze on the correct option, broken up into five quintiles. The quintiles were constructed by taking the difference between the proportional gaze for the correct (higher valued) and incorrect (lower valued) symbols on each trial, sorting the difference scores, and dividing them into five equal-sized bins, separately for each participant. The higher the quintile, the longer the correct option was fixated relative to the incorrect option. Error bars represent *±*1 standard error.

**Figure S15:**
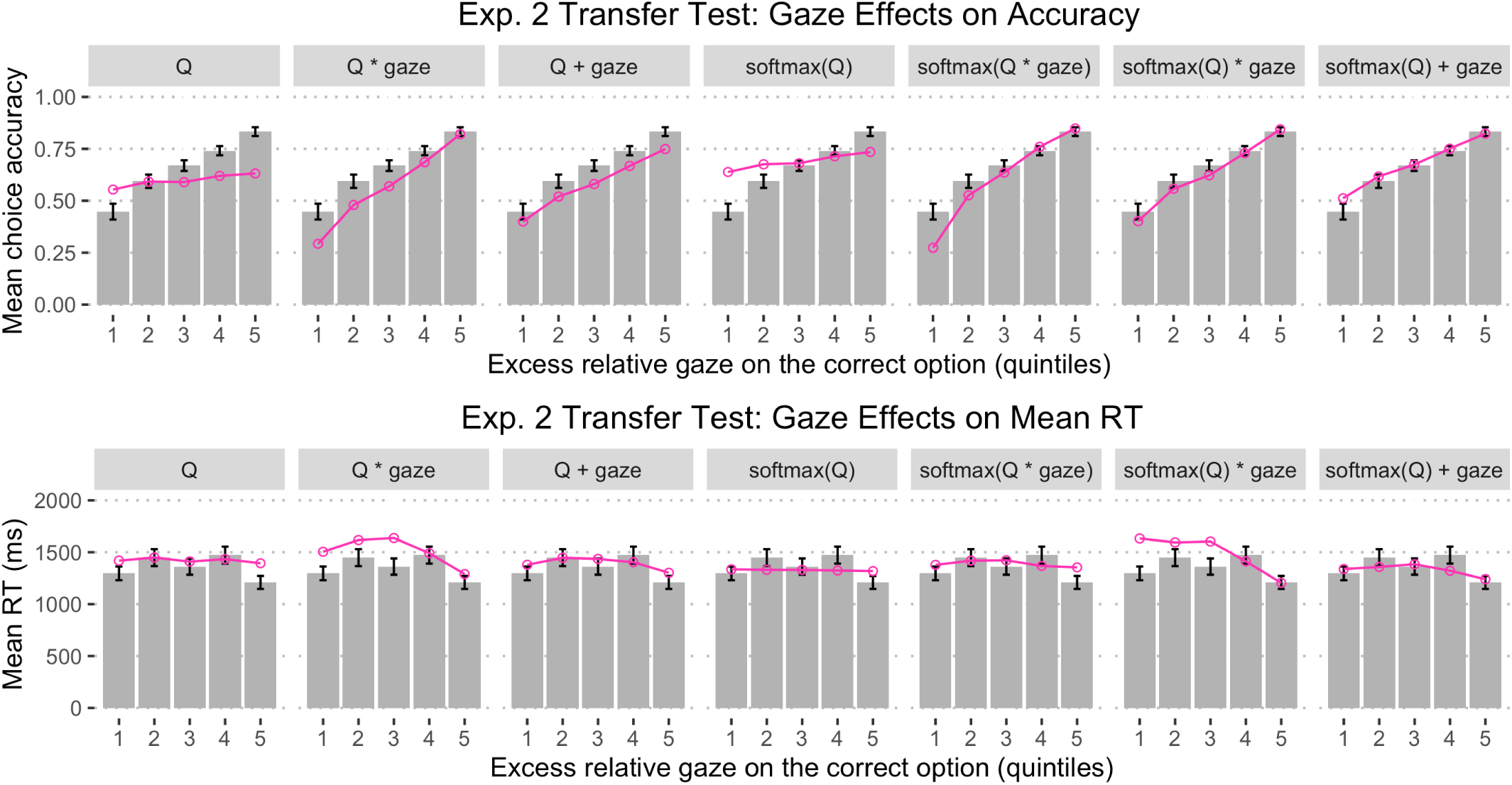
Model fit in Experiment 2: Mean choice accuracy (top) and mean RT (bottom) in the transfer test as a function of the excess proportional gaze on the correct option, broken up into five quintiles. The quintiles were constructed by taking the difference between the proportional gaze for the correct (higher valued) and incorrect (lower valued) symbols on each trial, sorting the difference scores, and dividing them into five equal-sized bins, separately for each participant. The higher the quintile, the longer the correct option was fixated relative to the incorrect option. Error bars represent *±*1 standard error.

**Table S1:** Multiple Regression Predicting Individual Choice Accuracy from RL-SSM Parameters (Exp 1)

| Predictor | b | SE | t | p |
| --- | --- | --- | --- | --- |
| Intercept | 0.76 | 0.13 | 6.04 | < .001 |
| Learning rate ($\alpha$) | 0.76 | 0.43 | 1.78 | .08 |
| Q drift scaling ($\beta_Q$) | 0.16 | 0.12 | 1.31 | .20 |
| Gaze drift scaling ($\beta_{gaze}$) | -0.18 | 0.12 | -1.48 | .14 |
| Start point upper bound ($A$) | -0.00011 | 0.00011 | -1.02 | .31 |
| Decision threshold ($b$) | 0.00016 | 0.000073 | 2.18 | .03 |
| Non-decision time ($t_0$) | -0.0011 | 0.0009 | -1.25 | .21 |
| *Adjusted $R^2 = .057, F(6, 76) = 1.82, p = .11$* |  |  |  |  |

**Table S2:** Multiple Regression Predicting Individual Choice Accuracy from RL-SSM Parameters (Exp 2: Learning Phase)

| Predictor | b | SE | t | p |
| --- | --- | --- | --- | --- |
| Intercept | 0.67 | 0.093 | 7.24 | < .001 |
| Learning rate ($\alpha$) | 0.20 | 0.091 | 2.24 | .03 |
| Relative encoding ($w_{rel}$) | 0.07 | 0.043 | 1.65 | .11 |
| Q drift scaling ($\beta_Q$) | 0.089 | 0.23 | 0.40 | .69 |
| Gaze drift scaling ($\beta_{gaze}$) | -0.25 | 0.22 | -1.15 | .26 |
| Start point upper bound ($A$) | -0.0017 | 0.00042 | -4.12 | < .001 |
| Decision threshold ($b$) | 0.0013 | 0.00037 | 3.60 | < .001 |
| Non-decision time ($t_0$) | 0.00028 | 0.00036 | 0.78 | .44 |
| *Adjusted $R^2 = .29$, $F(7, 42) = 3.80$, $p = .003$* |  |  |  |  |

**Table S3:** Multiple Regression Predicting Individual Mean RT from RL-SSM Parameters (Exp 1)

| Predictor | b | SE | t | p |
| --- | --- | --- | --- | --- |
| Intercept | 1714.77 | 292.43 | 5.86 | <.001 |
| Learning rate ($\alpha$) | 218.70 | 989.26 | 0.22 | .83 |
| Q drift scaling ($\beta_Q$) | -3526.42 | 282.13 | -12.50 | <.001 |
| Gaze drift scaling ($\beta_{gaze}$) | -3440.49 | 279.61 | -12.31 | <.001 |
| Start point upper bound ($A$) | -1.75 | 0.26 | -6.85 | <.001 |
| Decision threshold ($b$) | 3.33 | 0.17 | 19.78 | <.001 |
| Non-decision time ($t_0$) | 1.56 | 2.08 | 0.75 | .46 |
| *Adjusted $R^2 = .92$, $F(6, 76) = 160.8$, $p < .001$* |  |  |  |  |

**Table S4:**
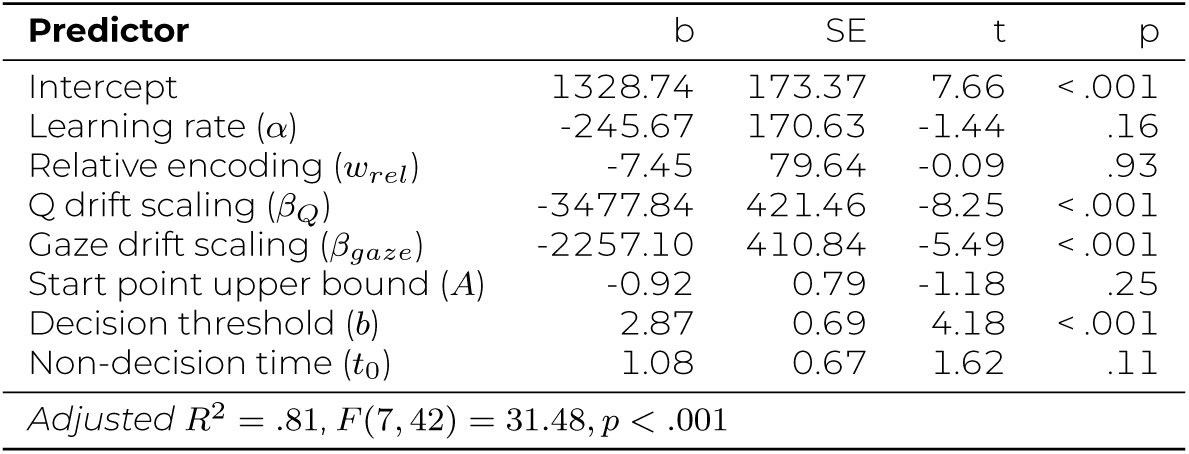
Multiple Regression Predicting Individual Mean RT from RL-SSM Parameters (Exp 2: Learning Phase)

**Table S5:** Multiple Regression Predicting Incongruent Trial Accuracy from RL-SSM Parameters (Exp 2: Transfer Test)

| Predictor | b | SE | t | p |
| --- | --- | --- | --- | --- |
| Intercept | 0.97 | 0.16 | 6.14 | < .001 |
| Learning rate ($\alpha$) | -0.08 | 0.16 | -0.53 | .60 |
| Relative encoding ($w_{rel}$) | -0.76 | 0.07 | -10.39 | < .001 |
| Q drift scaling ($\beta_Q$) | -0.85 | 0.38 | -2.22 | .03 |
| Gaze drift scaling ($\beta_{gaze}$) | -0.46 | 0.38 | -1.22 | .23 |
| Start point upper bound ($A$) | -0.0016 | 0.00072 | -2.23 | .03 |
| Decision threshold ($b$) | 0.0013 | 0.00063 | 2.09 | .04 |
| Non-decision time ($t_0$) | 0.00024 | 0.00061 | 0.39 | .70 |
| *Adjusted $R^2 = .72$, $F(7, 42) = 18.57$, $p < .001$* |  |  |  |  |

**Table S6:** Linear Mixed-Effects Model Predicting Proportional Gaze Advantage for the Correct Option from Trial Number, EV Difference and Overall Expected Value in Experiment 1.

| **Fixed Effects** | b | SE | t | p |
| --- | --- | --- | --- | --- |
| Intercept | 0.059 | 0.0062 | 9.44 | < .001 |
| EV Difference | 0.035 | 0.0052 | 6.60 | < .001 |
| Trial Number | 0.018 | 0.0052 | 3.44 | < .001 |
| Overall EV | 0.013 | 0.0070 | 1.89 | 0.062 |
| Trial Number × EV Difference | -0.0071 | 0.0051 | -1.37 | 0.17 |
| Trial Number × Overall EV | 0.0080 | 0.0050 | 1.60 | 0.11 |
| **Random Effects** | Variance |  |  |  |
| Intercept | 0.0014 |  |  |  |
| EV Difference | 0.00043 |  |  |  |
| Trial Number | 0.00035 |  |  |  |
| Overall EV | 0.0022 |  |  |  |
| Trial Number × EV Difference | 0.00031 |  |  |  |
| Trial Number × Overall EV | 0.00016 |  |  |  |
| Residual | 0.11 |  |  |  |
| *Improvement over intercept-only model: $\chi^2(25) = 83.61, p < .001$* |  |  |  |  |

**Table S7:** Linear Mixed-Effects Model Predicting Proportional Gaze Advantage for the Correct Option from Trial Number and Overall Expected Value in the Learning Phase of Experiment 2.

| **Fixed Effects** | b | SE | t | p |
| --- | --- | --- | --- | --- |
| Intercept | 0.20 | 0.015 | 12.65 | < .001 |
| Trial Number | 0.058 | 0.0093 | 6.20 | < .001 |
| Overall EV | 0.0026 | 0.0083 | 0.31 | 0.76 |
| Trial Number × Overall EV | -0.0067 | 0.0084 | -0.79 | 0.43 |
| **Random Effects** | Variance |  |  |  |
| Intercept | 0.010 |  |  |  |
| Trial Number | 0.0023 |  |  |  |
| Overall EV | 0.0015 |  |  |  |
| Trial Number × Overall EV | 0.0015 |  |  |  |
| Residual | 0.24 |  |  |  |
| *Improvement over intercept-only model: $\chi^2(12) = 122.8, p < .001$* |  |  |  |  |

**Table S8:** Logistic Mixed-Effects Model Predicting Choice Accuracy from Trial Number, EV Difference, Overall EV, and Proportional Gaze Advantage for the Correct Option in Experiment 1.

| **Fixed Effects** | b | SE | z | p |
| --- | --- | --- | --- | --- |
| Intercept | 1.63 | 0.13 | 12.28 | <.001 |
| Trial Number | 0.71 | 0.077 | 9.19 | <.001 |
| EV Difference | 0.67 | 0.072 | 9.20 | <.001 |
| Overall EV | -0.0091 | 0.052 | -0.17 | 0.862 |
| Gaze Difference | 1.076 | 0.074 | 14.58 | <.001 |
| Trial Number × EV Difference | 0.32 | 0.067 | 4.77 | <.001 |
| Trial Number × Overall EV | 0.019 | 0.052 | 0.37 | 0.71 |
| Trial Number × Gaze Difference | -0.14 | 0.060 | -2.29 | 0.022 |
| **Random Effects** | Variance |  |  |  |
| Intercept | 1.13 |  |  |  |
| Trial Number | 0.21 |  |  |  |
| EV Difference | 0.17 |  |  |  |
| Overall EV | 0.076 |  |  |  |
| Gaze Difference | 0.21 |  |  |  |
| Trial Number × EV Difference | 0.14 |  |  |  |
| Trial Number × Overall EV | 0.072 |  |  |  |
| Trial Number × Gaze Difference | 0.053 |  |  |  |
| *Improvement over no-gaze model: $\chi^2(17) = 708.15, p < .001$* |  |  |  |  |

**Table S9:** Logistic Mixed-Effects Model Predicting Choice Accuracy from Trial Number, Overall EV, and Proportional Gaze Advantage for the Correct Option in the Learning Phase of Experiment 2.

| **Fixed Effects** | b | SE | z | p |
| --- | --- | --- | --- | --- |
| Intercept | 2.59 | 0.19 | 13.54 | < .001 |
| Trial Number | 0.64 | 0.11 | 5.80 | < .001 |
| Overall EV | -0.03 | 0.079 | -0.38 | 0.71 |
| Gaze Difference | 1.33 | 0.097 | 13.66 | < .001 |
| Trial Number × Overall EV | 0.030 | 0.061 | 0.49 | 0.63 |
| Trial Number × Gaze Difference | -0.058 | 0.071 | -0.82 | 0.41 |
| **Random Effects** | Variance |  |  |  |
| Intercept | 1.49 |  |  |  |
| Trial Number | 0.34 |  |  |  |
| Overall EV | 0.13 |  |  |  |
| Gaze Difference | 0.25 |  |  |  |
| Trial Number × Overall EV | 0.033 |  |  |  |
| Trial Number × Gaze Difference | 0.042 |  |  |  |
| *Improvement over no-gaze model: $\chi^2(13) = 852.3, p < .001$* |  |  |  |  |

**Table S10:** Linear Mixed-Effects Model Predicting log RT from Trial Number, EV Difference, Overall EV, and Proportional Gaze Advantage for the Correct Option in Experiment 1.

| Fixed Effects | b | SE | t | p |
| --- | --- | --- | --- | --- |
| Intercept | 7.68 | 0.042 | 180.75 | < .001 |
| Trial Number | -0.070 | 0.017 | -4.11 | < .001 |
| EV Difference | -0.053 | 0.0067 | -7.84 | < .001 |
| Overall EV | -0.036 | 0.0082 | -4.43 | < .001 |
| Gaze Difference | -0.031 | 0.0068 | -4.61 | < .001 |
| Trial Number x EV Difference | -0.0089 | 0.0053 | -1.68 | 0.093 |
| Trial Number x Overall EV | -0.019 | 0.0059 | -3.16 | 0.0022 |
| Trial Number x Gaze Difference | -0.0097 | 0.0052 | -1.85 | 0.065 |
| Random Effects | Variance |  |  |  |
| Intercept | 0.15 |  |  |  |
| Trial Number | 0.022 |  |  |  |
| EV Difference | 0.0015 |  |  |  |
| Overall EV | 0.0034 |  |  |  |
| Gaze Difference | 0.0015 |  |  |  |
| Trial Number x EV Difference | 0 |  |  |  |
| Trial Number x Overall EV | 0.00061 |  |  |  |
| Trial Number x Gaze Difference | 0 |  |  |  |
| Residual | 0.13 |  |  |  |
| Improvement over no-gaze model: $\chi^2(4) = 31.45, p < .001$ | | | | |
| Note: Random effect correlations excluded from the model to aid convergence. |  |  |  |  |

**Table S11:** Linear Mixed-Effects Model Predicting log RT from Trial Number, Overall EV, and Proportional Gaze Advantage for the Correct Option in the Learning Phase of Experiment 2.

| **Fixed Effects** | b | SE | t | p |
| --- | --- | --- | --- | --- |
| Intercept | 7.03 | 0.045 | 155.40 | < .001 |
| Trial Number | -0.19 | 0.015 | -13.10 | < .001 |
| Overall EV | -0.0099 | 0.010 | -0.99 | 0.33 |
| Gaze Difference | -0.071 | 0.0091 | -7.72 | < .001 |
| Trial Number × Overall EV | 0.0031 | 0.0072 | 0.42 | 0.67 |
| Trial Number × Gaze Difference | 0.027 | 0.0080 | 3.32 | .0018 |
| **Random Effects** | Variance |  |  |  |
| Intercept | 0.101 |  |  |  |
| Trial Number | 0.0091 |  |  |  |
| Overall EV | 0.0034 |  |  |  |
| Gaze Difference | 0.0025 |  |  |  |
| Trial Number × Overall EV | 0.00097 |  |  |  |
| Trial Number × Gaze Difference | 0.0015 |  |  |  |
| Residual | 0.18 |  |  |  |
| *Improvement over no-gaze model: $\chi^2(13) = 184.7, p < .001$* |  |  |  |  |

**Table S12:** Logistic Mixed-Effects Model Predicting Choice Accuracy from EV Difference, Relative Value Difference, Overall EV, Overall Relative Value, and Proportional Gaze Advantage for the Correct Option in the Transfer Test of Experiment 2.

| **Fixed Effects** | b | SE | z | p |
| --- | --- | --- | --- | --- |
| Intercept | 1.23 | 0.21 | 5.78 | < .001 |
| EV Difference | 0.27 | 0.10 | 2.57 | .010 |
| Relative Value Difference | 1.20 | 0.16 | 7.62 | < .001 |
| Overall EV | -0.16 | 0.11 | -1.41 | 0.16 |
| Overall Relative Value | 0.0016 | 0.065 | 0.025 | 0.98 |
| Gaze Difference | 0.89 | 0.11 | 7.75 | < .001 |
| **Random Effects** | Variance |  |  |  |
| Intercept | 2.088 |  |  |  |
| EV Difference | 0.40 |  |  |  |
| Relative Value Difference | 1.08 |  |  |  |
| Overall EV | 0.55 |  |  |  |
| Overall Relative Value | 0.12 |  |  |  |
| Gaze Difference | 0.52 |  |  |  |
| *Improvement over no-gaze model: $\chi^2(7) = 540.57, p < .001$* |  |  |  |  |

**Table S13:** Linear Mixed-Effects Model Predicting log RT from EV Difference, Unsigned Relative Value Difference, Unsigned Proportional Gaze Difference, Overall Expected Value and Overall Relative Value in the Transfer Test of Experiment 2.

| **Fixed Effects** | b | SE | t | p |
| --- | --- | --- | --- | --- |
| Intercept | 7.024 | 0.051 | 137.41 | < .001 |
| EV Difference | -0.0030 | 0.0099 | -0.31 | 0.76 |
| Unsigned Relative Value Difference | -0.049 | 0.011 | -4.63 | < .001 |
| Unsigned Gaze Difference | -0.080 | 0.013 | -5.95 | < .001 |
| Overall EV | -0.011 | 0.012 | -0.96 | 0.34 |
| Overall Relative Value | -0.054 | 0.0084 | -6.49 | < .001 |
| **Random Effects** | Variance |  |  |  |
| Intercept | 0.13 |  |  |  |
| EV Difference | 0.0030 |  |  |  |
| Unsigned Relative Value Difference | 0.0038 |  |  |  |
| Unsigned Gaze Difference | 0.0072 |  |  |  |
| Overall EV | 0.0051 |  |  |  |
| Overall Relative Value | 0.0016 |  |  |  |
| Residual | 0.20 |  |  |  |
| *Improvement over no-gaze model: $\chi^2(7) = 281.76, p < .001$* |  |  |  |  |

